# *De novo* Autogenic Engineered Living Functional Materials

**DOI:** 10.1101/2025.01.21.634218

**Authors:** Hoda M. Hammad, Seth Swarnadeep, Erin R. Crater, Robert B. Moore, Sanket Deshmukh, Avinash Manjula-Basavanna, Anna M. Duraj-Thatte

## Abstract

Autogenic engineered living materials (ELMs) involve *in situ* production and engineering of native extracellular matrix (ECM). However, the existing autogenic ELMs have limited scope and functionalities. Herein, we report a platform for *de novo* autogenic functional ELMs. By protein mining, we have discovered CsgA-like 33,564 homologs that can find potential utility as *de novo* ECM of protein nanofibers. By employing AlphaFold2 and molecular dynamics simulations, we shed insights into the CsgA-like β-solenoid protein structures and stability. By hacking *Escherichia coli* curli machinery, we demonstrate the production of *de novo* autogenic ELMs from CsgA-like proteins (≤9-times the molecular weight and β-sheet repeat units) of extremophilic non-model bacteria. Additionally, we biomanufacture macroscopic materials with tunable mechanical properties (enhancing storage modulus by 3-times) and programmable functionalities (3D printability, binding to nanoparticles/antibodies). This work showcases a versatile platform to discover, rationally design, and harness the sophisticated functionalities of natural systems for futuristic autogenic ELMs.

---

Engineered living materials (ELMs) is a rapidly growing field wherein living cells are engineered to produce materials with life-like functionalities for sustainability, biomedicine, biosensing, biomining, and bioremediation applications.^1–8^ Based on the polymeric matrix, ELMs can be broadly classified into two types, namely, exogenic and autogenic ELMs.^9, 10^ In exogenic ELMs, living cells are embedded in a synthetic/natural polymeric matrix obtained from an external source, whereas, in autogenic ELMs, the living cells are utilized/engineered for *in situ* production of native/functional polymeric matrix.^11^ Evidently, autogenic ELMs are advantageous over exogenic ELMs, as they enable on-demand *in situ* biosynthesis, self-assembly, self-organization, self-regeneration, self-regulation, environmental adaptability, and functionalization of the polymeric matrix.^12–14^ Although the autogenic ELMs are crucial to realize the full potential of this emerging technology, there are only a few reported autogenic ELMs, emphasizing the need for innovative design strategies to 1) harness the programmability and biomanufacturing capabilities of living cells and 2) discover the wide variety of functionalities prevalent in the natural systems.^15–19^

In the existing autogenic ELMs, the native extracellular matrix (ECM) of bacterial biofilms is utilized either directly or after functionalization. For example, the native protein nanofibers-based ECM of *Escherichia coli* and *Bacillus subtilis*, which comprises CsgA and TasA, respectively, have been genetically modified by fusing them with the desired protein domains to obtain functional autogenic ELMs.^15, 20–22^ Alternatively, the native cellulose nanofibers-based ECM of *Komagataeibacter rhaeticus* has been employed to obtain autogenic ELMs, but unlike the above example of protein-based ECM, the genetic functionalization of cellulose/polysaccharide to controllably modulate its properties is still elusive.^6, 23^ On the other hand, the genetic modification of surface-layer proteins of *E. coli*, *B. subtilis,* and *Caulobacter crescentus* has resulted in autogenic ELMs, but they have limited scope to tailor the physical, chemical, and biological properties in comparison to functional nanofibers-based ECM.^12, 24, 25^

In the last twelve years, the ECM protein nanofibers of *E. coli* self-assembled from CsgA are the most studied autogenic ELMs.^7, 26^ These protein nanofibers, commonly known as curli, are attractive due to their 1) resistance to heat, solvents, pH, detergents, and denaturants, 2) mechanical properties, and 3) ability to modulate functionalities by genetically grafting the desired protein domains.^27, 28^ CsgA, with its cross-β structure, is energetically among the most stable protein folds, while the β-solenoid architecture facilitates the positioning of the C- and N-terminal on its periphery. As a result, the peptide/protein domains genetically grafted to CsgA via a flexible linker are often found to fold independently to their functional form (**Supplementary Fig. 1**).^26^ Moreover, the head-to-tail stacking of CsgA building blocks leads to the formation of curli nanofibers, and the genetically grafted peptide/protein domain gets displayed on its periphery to form the functional curli nanofibers, which have been demonstrated for various applications in the last several years.^29, 30^ Thus, we hypothesized that the β-solenoid architecture of the CsgA protein building block is crucial for the fascinating properties and applications of curli nanofibers-based autogenic ELMs.

In this work, we present *de novo* autogenic functional ELMs, wherein the *E. coli* is engineered to produce *de novo* ECM of protein nanofibers (**Fig. 1**). Motivated by the intriguing characteristics described above, we conducted protein database mining to discover CsgA-like β-solenoid proteins that can serve as *de novo* building blocks of synthetic ECM. We utilized the artificial intelligence (AI) tool AlphaFold2 to predict the structure of these *de novo* β-solenoid proteins, and the stability of the folded structure was deciphered using all-atom (AA) molecular dynamics (MD) simulations in the explicit water model.^31^ As these β-solenoid proteins were from non-model organisms thriving in different parts of the globe, we have hacked the curli secretion machinery of model organism *E. coli* to produce *de novo* ECM. Further, we employ these ECMs to produce functional materials like hydrogels/films and show that their physicochemical properties can be tailored. In addition, we demonstrate that the biological functions of these *de novo* ECM-based ELMs could be controllably modified by genetically grafting the desired protein domains to facilitate specific binding to nanoparticles or antibodies.

**Figure 1.**
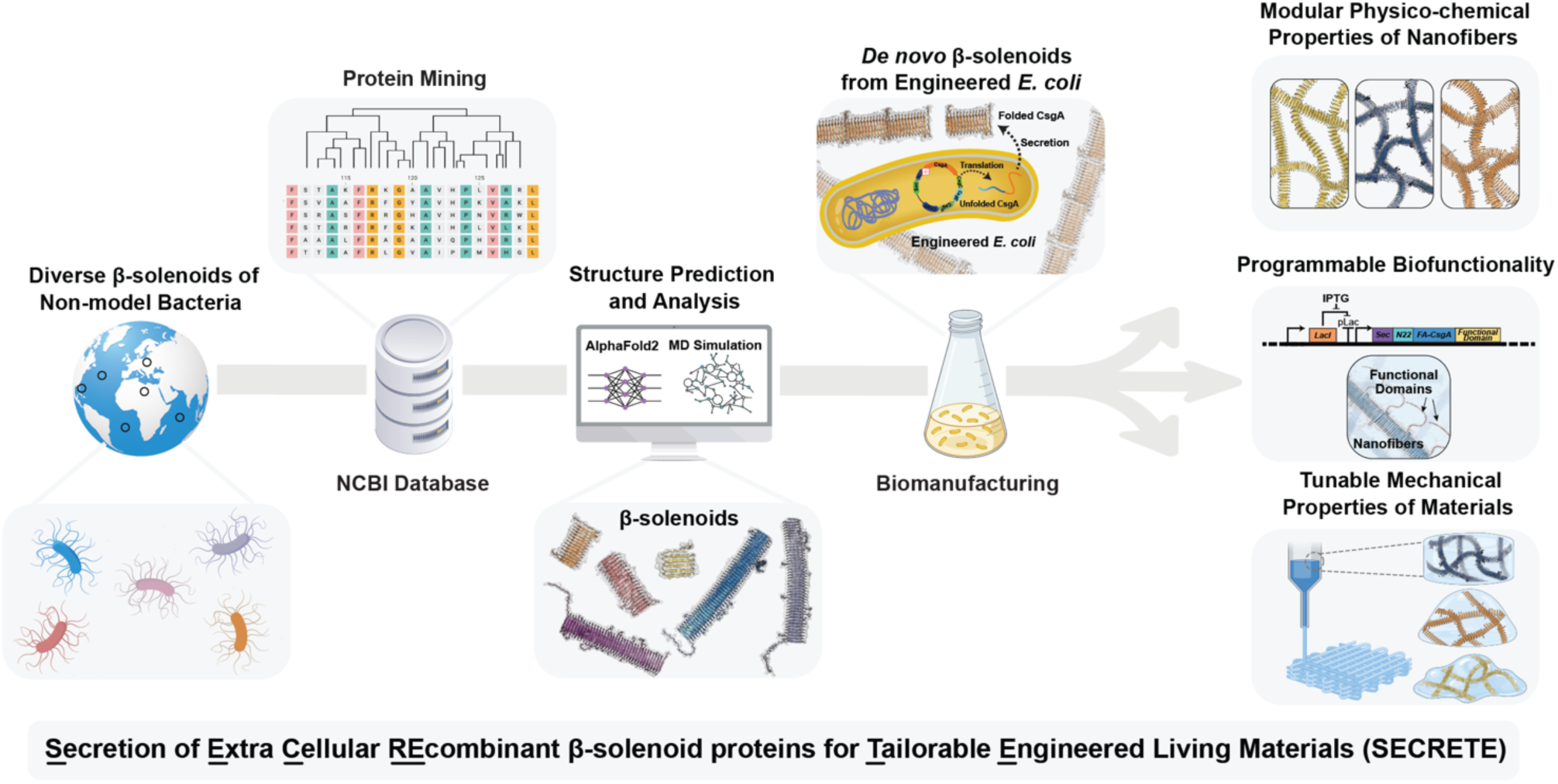
Schematic of the SECRETE platform to produce CsgA-like *de novo* β-solenoid proteins for autogenic engineered living functional materials. CsgA-like proteins discovered by mining the National Center for Biotechnology Information (NCBI) protein database were studied by using AlphaFold2 and molecular dynamics (MD) simulations to predict their structure. *E. coli* curli machinery was hacked to produce *de novo* β-solenoid proteins of non-model extremophilic bacteria to obtain materials with modular physico-chemical properties, programmable biofunctionalities, and tunable mechanical properties.

## Results

### Protein mining and structure prediction

CsgA is a 13 kDa protein of 131 amino acids with five cross-β repeats, wherein the minimalistic curli repeat sequence is presented in the form X_6_QXGX_2_NX_10_.^32, 33^ To identify CsgA-like proteins, we utilized the National Center for Biotechnology Information (NCBI) database to search for proteins having X_6_QXGX_2_NX_10_ sequence, which resulted in 33,564 entries and their lengths varied from 141 to 1390 amino acids. From these, we randomly selected 50 protein sequences, and their protein structure was predicted using AlphaFold2, which revealed β-solenoid architecture for all of them (**Supplementary Fig. 2**). Finally, for a detailed study, we selected five protein sequences of different lengths that belonged to diverse bacteria species, namely, *Halomonas saliphila* (Hs13-CsgA), *Alteromonas macleodii* (Am18-CsgA), *Blastomonas sp. CACIA14H2* (Bc36-CsgA), *Erythrobacter longus* (El43-CsgA), and *Ensifer sp. Root31* (Er46-CsgA). These CsgA homologs, named according to their genus, species, and predicted number of cross-β repeats, represent a range of ecological adaptations and functional properties, none of which had been experimentally validated for curli production to the best of our knowledge, prior to this study (**Supplementary Fig. 3**).

The 3D structure of CsgA and five other CsgA homologs predicted using AlphaFold2 revealed the signature sequence KR_1_-Ω-Ψ-Ω-Ψ-Ω-KR_7_ of cross-β sheet structure, wherein Ω represents outward-facing residues, while KR_1_, KR_7,_ and Ψ represent inward-facing residues (**Fig 2. and Supplementary Fig. 4**). Each repeat unit in CsgA consists of two antiparallel β-strands, β_1,_ and β_2_, arranged in a cross-β sheet structure, with each β-strand made of seven residues from KR_1_ to KR_7_ that align across the entire folded monomer. These β-strands within each repeat unit are connected by short loops. The first loop, connecting β_1_ to β_2_ within a repeat unit, is referred to as the intra-repeat loop, while the loop connecting one repeat unit to the next is called the inter-repeat loop. Both loops are typically composed of four residues, while the intra-repeat loop is represented by the sequence X-G-X-G, where X represents variable residues, and G signifies the conserved glycine, which allows for structural flexibility. In summary, the conserved β-strand and loop sequence of each repeat unit follows the pattern: β_1_ strand (KR_1_-Ω-Ψ-Ω-Ψ-Ω-KR_7_), intra-repeat loop (X-G-X-G), β_2_ strand (KR_1_-Ω-Ψ-Ω-Ψ-Ω-KR_7_), inter-repeat loop (X-X-X-X), and so on. Different CsgA homologs vary in the number of repeat units: *E. coli* CsgA consists of 5 repeat units, while Hs13-CsgA has 13, Am18-CsgA has 18, Bc36-CsgA has 36, El43-CsgA has 43, and Er46-CsgA has 46 repeat units (43 in the core β-solenoid structure and 3 in the outlier region).

**Figure 2.**
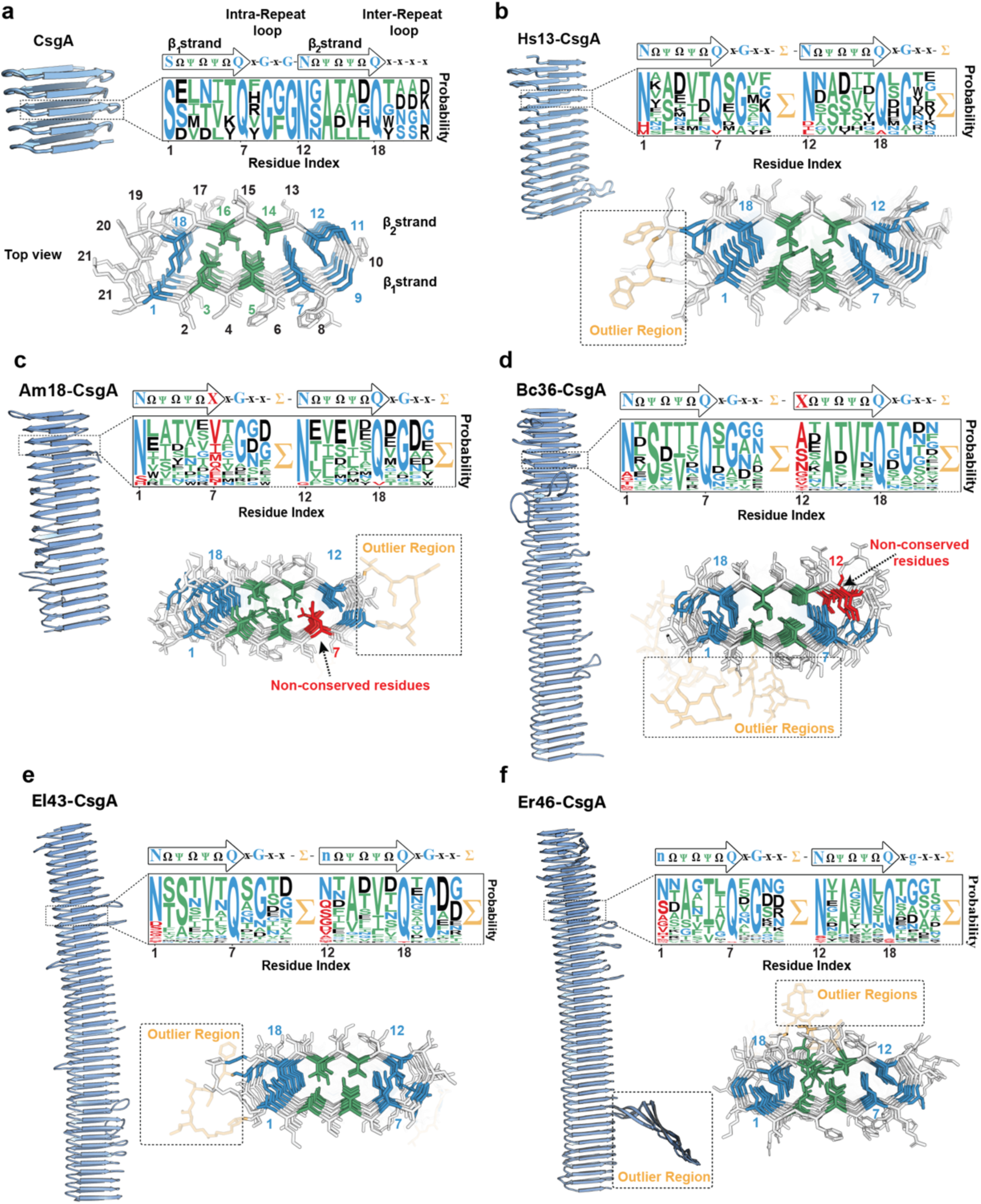
Structures of β-solenoid protein building blocks predicted by AlphaFold2. The predicted β-solenoid protein models of **a** CsgA, **b** Hs13-CsgA, **c** Am18-CsgA, **d** Bc36-CsgA, **e** El43-CsgA, and **f** Er46-CsgA. The longitudinal view of β-solenoid protein model shows the number of cross-β repeats. Each β-solenoid protein sequence is presented to showcase the conserved (N, Q, G indicates >50% of conservation probability) and non-conserved residues (n, g indicates 25-50% of conservation probability and for <25%, it is indicated as X). On-axis top view shows the β-solenoid fold with inward-facing residues (green), key residues (blue), non-conserved residues (red), and outlier regions (orange). For clarity, only the top 5 cross-β repeats are shown for the on-axis top view.

Within each β-strand pair (β_1_ and β_2_) in a repeat unit, key residues (KR), located at the beginning (KR_1_) and the end (KR_7_) of each β-strand, are reported to play a crucial role in stabilizing and ensuring proper folding of the monomer.^32, 33^ In *E. coli* CsgA, KR_1_ of β_1_ is conserved as serine (S), while KR_1_ of β_2_ is asparagine (N), and KR_7_ of both β_1_ and β_2_ is glutamine (Q).^34^ However, these key residues KR_1_ / KR_7_ in β_1_ / β_2_ strands are conserved to various extents for the other five β-solenoids, and the non-conserved key residues, denoted as X in the consensus motif, are highlighted in red. The inward-facing residues, Ψ, shown in green, are tightly packed between the β-sheets, stabilizing the fibril core. These inward-facing residues typically consist of hydrophobic amino acids (A, I, V, L, F), as well as serine (S) or threonine (T). Meanwhile, outward-facing residues, Ω, shown in black, are usually hydrophilic and exposed to the surrounding environment, facilitating fibrillation. Additionally, our structural analysis revealed outlier regions (highlighted in orange and denoted as ∑), which extend beyond the canonical β-strands and loop motifs, are observed in all five variants, except for *E. coli* CsgA. While the precise role of these outlier regions remains unclear, they may have functional implications or impact the overall architecture of the fibrils, warranting further investigation.

Additional analysis shed insights into the physicochemical characteristics of these six β-solenoid proteins (**Fig. 3a-d, Supplementary Table 1-2**). The molecular weight of the proteins increased with a greater number of repeats, from 13.1 kDa for CsgA to 117.6 kDa for Er46-CsgA, while their isoelectric points remained similar, ranging from 3.3 to 4.5. The predicted charge of CsgA at pH 7.4 was −6.3, and it increased by several folds for the five other β-solenoid proteins, with the highest charge of −116.6 observed for El43-CsgA. The hydrophobicity, as predicted by their Gravy score (−0.7 to −0.1) indicated that all five *de novo* β-solenoid proteins are more hydrophobic than CsgA (**Supplementary Fig. 5**).

**Figure 3.**
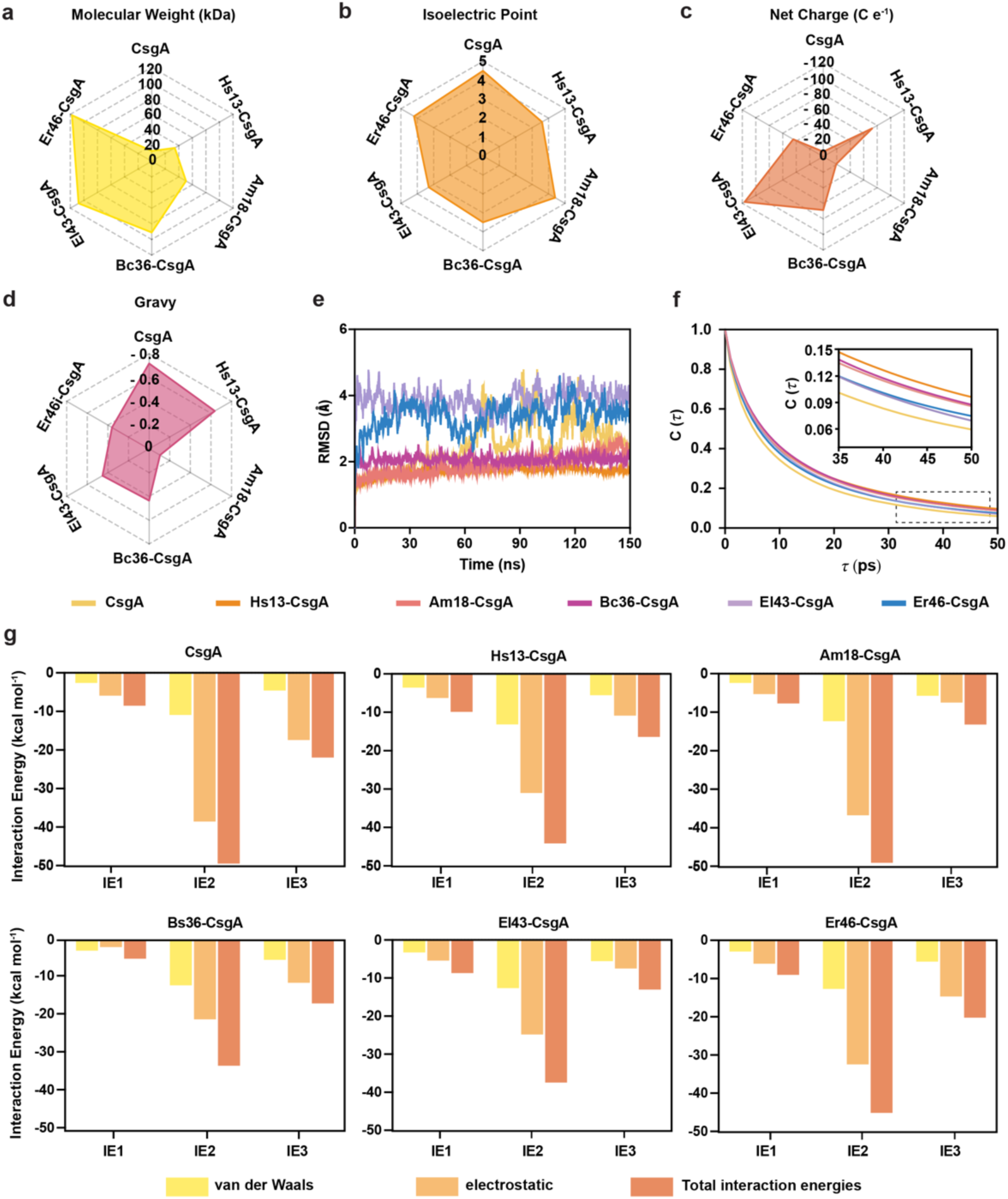
Physicochemical characteristics of β-solenoid proteins and their structure stability analysis by all-atom molecular dynamics simulations. Comparison of the physicochemical characteristics of β-solenoid proteins, **a** molecular weight, **b** isoelectric point, **c** net charge, and **d** hydrophobicity index. **e** Root mean square deviation (RMSD) of the β-strands for β-solenoid proteins throughout MD simulation. **f** Hydrogen bond autocorrelation function for β-solenoid proteins. **g** Average normalized van der Waals, electrostatic, and total interaction energies (IE). IE1: between KR and KR, IE2: KR and all residues, and IE3: non-KR and all residues, over the period of the production run. The energies were normalized per residue energies.

### Decoding β-solenoid protein structure stability

All six β-solenoid proteins showed pLDDT (predicted local distance difference test) confidence scores between 71 and 89, which indicated a good backbone prediction (**Supplementary Fig. 6**). Using structures predicted by AlphaFold2, we conducted AA MD simulations to gain molecular-level insights into the stability of β-solenoid proteins, focusing on both structural and energetic analyses (**Supplementary Video 1**). For all six proteins, the root mean square deviation (RMSD) of all heavy atoms in backbone and the side chains of amino acids in β-sheets, increases with increasing residues (initial equilibration of less than ∼5 ns), and it remains stable during the remainder of the simulations run (**Fig. 3e**). RMSD of the entire protein showed slightly more perturbations and higher RMSD values, which could be due to the fluctuations in the N-terminal and outlier regions, which is consistent with the pLDDT scores (**Supplementary Fig. 7-8**). The residue-wise root mean squared fluctuations (RMSF) was consistent with the RMSD analysis, showing that the N-terminal and outlier residues had more fluctuations than the β-sheet region residues (**Supplementary Fig. 9**).

The Ramachandran plot showed a dense cluster of points with *ϕ* ≈ −120° and *ψ* ≈ 120° throughout 150 ns MD simulations, suggesting that the overall backbone dihedral angles are relatively preserved in the β-sheet region, with minor presence of α-sheets in all proteins (right α-sheet: *ϕ* = −60° and *ψ* = −45°; left α-sheet: *ϕ* = 70° and *ψ* = 35°; **Supplementary Table 3**, **Supplementary Fig. 10**).^35^ The hydrogen bond characteristics between all protein residue pairs forming hydrogen bonds were evaluated using a geometric criterion described in the Method section.^36^ The hydrogen bond analysis indicates that the presence of bifurcated hydrogen bonds, in addition to hydrogen bonds in β-sheets, may further contribute to the structural stability of the proteins.^37^ A slower decay of hydrogen bond autocorrelation function was observed for the five *de novo* β-solenoid proteins compared to CsgA, which indicates that the hydrogen bonds formed between amino acid pairs in the five *de novo* β-solenoid proteins are more stable compared to those in CsgA protein.

A detailed structural analysis revealed that the five β-solenoid proteins contain a higher number of charged residues (e.g., D, E, K, R) compared to CsgA (**Supplementary Fig. 11**). Additionally, the local environment of each residue (within 12 Å) is predominantly composed of glycine (G) and polar residues (e.g., N, Q, S, T) rather than non-polar residues (e.g., I, L, M, F, W) (**Supplementary Fig. 12**). To investigate further the role of charged residues and their local environments, we calculated averaged non-bonded interaction energies (IE) between KR and KR (IE1), between KR and all residues (IE2) and between non-KR and all residues (IE3) (**Fig. 3g**). Across all six proteins, the electrostatic interactions in IE1, IE2, and IE3 were generally more favorable (more negative) compared to van der Waals interactions but exhibited varying patterns. The electrostatic interactions in IE1 were consistent across most proteins, except for Bc36-CsgA, which showed less favorable interactions (less negative). In contrast, electrostatic interactions in IE2 and IE3 displayed greater variability, with CsgA and Er46-CsgA showing more favorable energies than the other proteins. Notably, IE2 electrostatic energies were more favorable than IE3. These results highlight that both charge placement and the local environment of these residues play an important role in determining electrostatic energies. Meanwhile, van-der Waals energy contributions in IE1, IE2, and IE3 were distinct but consistent across all six proteins, likely due to their similar geometries.^38^ Overall, MD simulations revealed that all six β-solenoid proteins are stabilized by hydrogen bond formation among the β-sheets and favorable electrostatic interactions between the key residues and all residues.

### Biomanufacturing of modular *de novo* autogenic engineered living materials

The above described five *de novo* CsgA homologs are from extremophilic non-model bacteria, which poses several challenges, such as culturing bacteria, scalable production, and genetic modifications. To circumvent these challenges, we have hacked the curli secretion machinery of *E. coli* to develop a platform termed **SECRETE**, which stands for **S**ecretion of **E**xtra **C**ellular **RE**combinant β-solenoid proteins for **T**ailorable **E**ngineered living materials. Herein, we employed *E. coli* strain PQN4, wherein the curli operon is deleted, which makes it suitable for curli overproduction. These *de novo* β-solenoid protein monomers were designed by integrating Sec (N-terminal signal sequence) and N22 (N-terminal curli-specific targeting sequence) sequences of *E. coli* to facilitate extracellular secretion and self-assembly.

Congo Red dye assay was utilized to experimentally and qualitatively assess the cross-β structure of Hs13-CsgA, Am18-CsgA, Bc36-CsgA, El43-CsgA, and Er46-CsgA (**Fig. 4a**). All these five CsgA homologs showed good CR-binding like that of CsgA; the relative differences in the Congo red binding could be attributed to slightly different 1) affinities due to their distinct amino acid compositions and structure and 2) levels of production of CsgA homologs. Wide-angle X-ray scattering (WAXS) analysis revealed characteristic *d*-spacing (interplanar spacing) values of 0.98 nm and 0.46 nm, corresponding to inter-β-sheet and inter-β-strand distances, respectively, consistent with that of CsgA, indicating the cross-β architecture of CsgA homologs (**Fig. 4b, c**).^39^ Further, ultrastructural characterization of bacterial cultures by field-emission scanning electron microscopy (FESEM) showed the protein nanofibers self-assembled in the extracellular milieu (**Fig. 4d**). Moreover, the characteristic birefringent property like CsgA was observed for all CsgA homolog nanofibers, under crossed polarizers (**Fig. 4e**). Subsequently, to demonstrate the biomanufacturing of macroscopic modular ELMs, we produced hydrogels of all six variants by using the filtration protocol^18^, and the FESEM images showed the dense network of nanofibers in the hydrogels (**Fig. 5a**). In addition, the hydrogels were cast on templates and dried under ambient conditions to fabricate aquaplastics, which could find potential applications as biodegradable bioplastics, robust coatings and sustainable materials (**Fig. 5b**). ^4^

**Figure 4.**
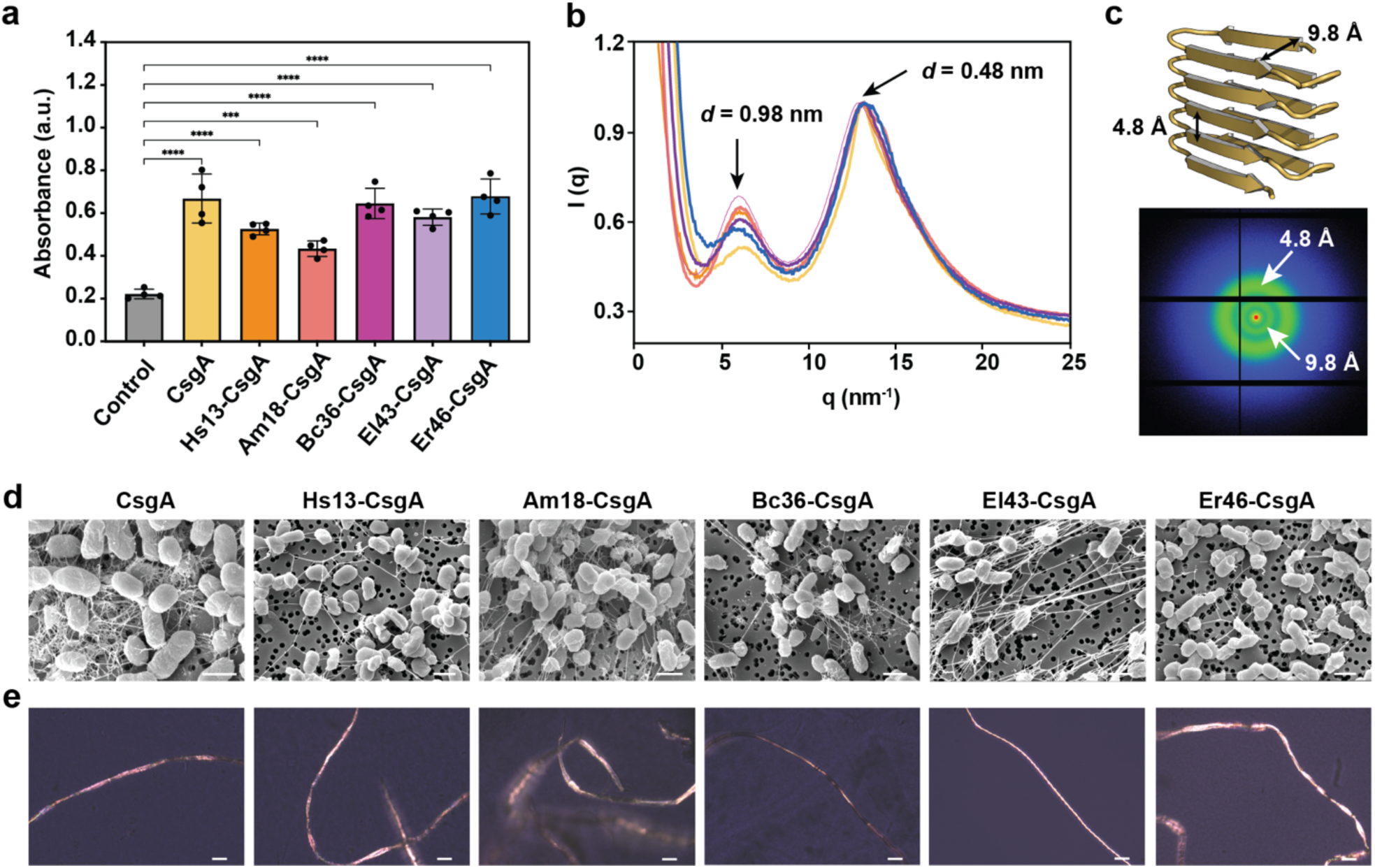
SECRETE platform to produce β-solenoid proteins of non-model organisms by hacking the curli secretion machinery of *E. coli*. a Congo red assay to determine the β-solenoid proteins produced by engineered *E. coli*. Biological replicates n = 4. Data represented as mean ± standard deviation. ****p ≤ 0.0001, one-way ANOVA followed by Dunnett’s test. **b** Wide-angle X-ray scattering (WAXS) analysis revealed cross-β characteristics of β-solenoid proteins. c. *d*-spacing (interplanar) values of 0.98 nm and 0.46 nm, corresponding to inter-β-sheet and intra-β-strand distances. **d.** Field-emission scanning electron microscopy (FESEM) images show the β-solenoid protein nanofibers self-assembled in the extracellular milieu. Scale bar 1 μm. **e.** Congo red birefringence showed the cross-β characteristics of β-solenoid protein nanofibers. Scale bar 50 μm.

**Figure 5.**
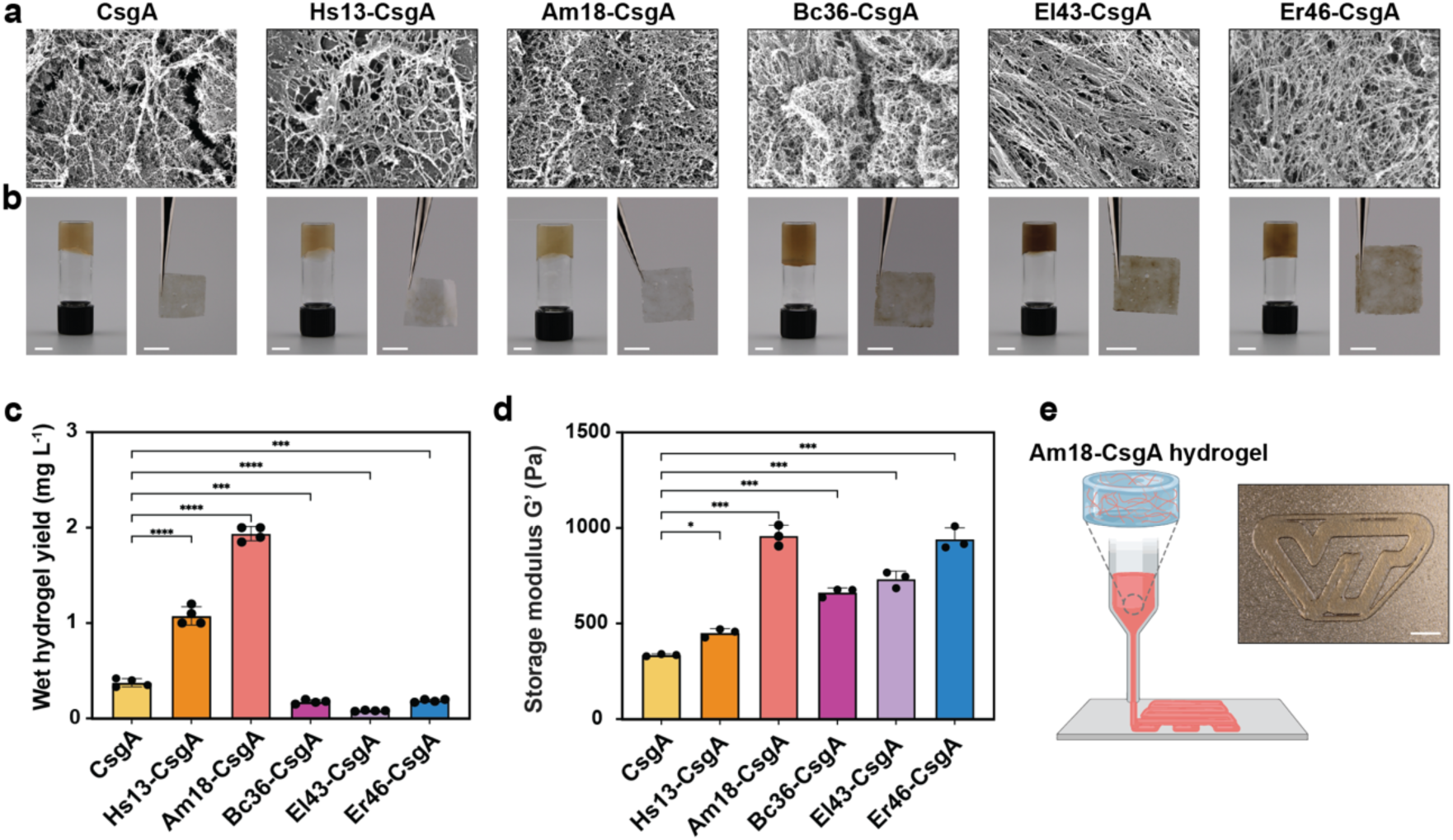
Biomanufacturing of macroscopic autogenic engineered living materials from various β-solenoid proteins. **a** Field-emission scanning electron microscopy (FESEM) images of β-solenoid protein hydrogels. Scale bar 1 μm. **b** Optical images of hydrogel and film manufactured using β-solenoid protein nanofibers. Scale bar 0.5 cm. **c** Yield of β-solenoid protein hydrogel obtained from engineered *E. coli* using filtration protocol. Biological replicates n = 4. Data represented as mean ± standard deviation. ****p ≤ 0.0001, ***p ≤ 0.0009 one-way ANOVA followed by Dunnett’s test. **d** Storage modulus of β-solenoid protein hydrogel. Biological replicates n = 3. Data represented as mean ± standard deviation. ***p ≤ 0.001, *p ≤ 0.014 one-way ANOVA followed by Dunnett’s test. **e** 3D printing of Am18-CsgA protein hydrogel. Scale bar 1 cm.

The yield of the hydrogels varied, Am18-CsgA (1938 ± 75 mg L^−1^) and Bs13-CsgA (1075 ± 96 mg L^−1^) showed the highest yields, both exceeding that of *E. coli* CsgA (375 ± 42 mg L^−1^), making them more suitable for scalable biomaterial production (**Fig. 5c**). As the five *de novo* β-solenoid proteins have more cross-β repeats than CsgA and the nanofibers network observed in the FESEM images were slightly different, we anticipated that their mechanical properties could be different. To test this, we investigated the storage modulus (G’) of hydrogels, which revealed that all five *de novo* β-solenoids exhibited significantly higher G’ values than CsgA. Interestingly, G’ was found to increase (by two folds) with the number of cross-β repeats (from 5 to 46). However, G’ of Am18-CsgA with 18 repeats was also the same as that of Er46-CsgA with 46 repeats, which could be attributed to differences in the 1) physicochemical properties such as greater hydrophobicity and lower charge, 2) aggregation of nanofibers and in turn the interactions with water that facilitates gelation (**Fig. 5d**). In addition, these curli hydrogels are suitable for extrusion bioprinting due to their shear-thinning behavior, a key property that enables it to flow like a fluid under the influence of external stress and upon removal of the external stress, it reverts to its initial viscosity. Building on this property, we demonstrated the 3D printability of *de novo* β-solenoid protein hydrogels by printing the Virginia Tech University logo using Am18-CsgA, which was utilized due to the higher yield of hydrogel (**Fig. 5e, Supplementary Fig. 13, Supplementary Video 2**).

Having demonstrated the modulation of physicochemical properties of *de novo* β-solenoid protein building block-based nanofibers and macroscopic materials, we next explored their potential for programming the biological functionalities. To achieve this, we selected Hs13-CsgA as the β-solenoid protein building block, which was genetically fused with a functional peptide or protein to assess whether the engineered variants could be secreted and self-assembled into biofunctional nanofibers. Herein, the C-terminal of Hs13-CsgA was genetically fused with 8-amino-acid iron-binding peptide (Hs13-CsgA-IronBP) or 185-amino-acid Immunoglobulin G (IgG) antibody binding protein^40^ (Hs13-CsgA-IgG-BD) via a 36-amino-acid flexible linker. AlphaFold2 structure predictions confirmed that both Hs13-CsgA-IronBP and Hs13-CsgA-IgG-BD retained their characteristic 13 repeats β-solenoid architecture with no structural disturbances caused by the fused peptide or protein domain (**Fig. 6, Supplementary Fig. 14**). Congo red assay and FESEM images of Hs13-CsgA-IronBP and Hs13-CsgA-IgG-BD confirmed their cross-β structure and nanofibrillar assemblies, respectively (**Supplementary Fig. 15**). The iron-binding assay showed that Hs13-CsgA-IronBP had a significantly higher weight of bound iron oxide nanoparticles than the wildtype Hs13-CsgA nanofibers. The elemental composition studied by using energy dispersive X-ray analysis (EDAX) also confirmed that Hs13-CsgA-IronBP has significantly higher iron content (**Fig. 6a, Supplementary Fig. 16**). Similarly, Hs13-CsgA-IgG-BD was found to bind red fluorescent IgG antibodies nearly twice higher than wildtype Hs13-CsgA nanofibers, as indicated by fluorescence intensity measurements and imaging (**Fig. 6b**). These results highlight the utility of *de novo* β-solenoid nanofibers to tailor their functionalities for biomining, bioremediation, biosensing, and biomedical applications.

**Figure 6.**
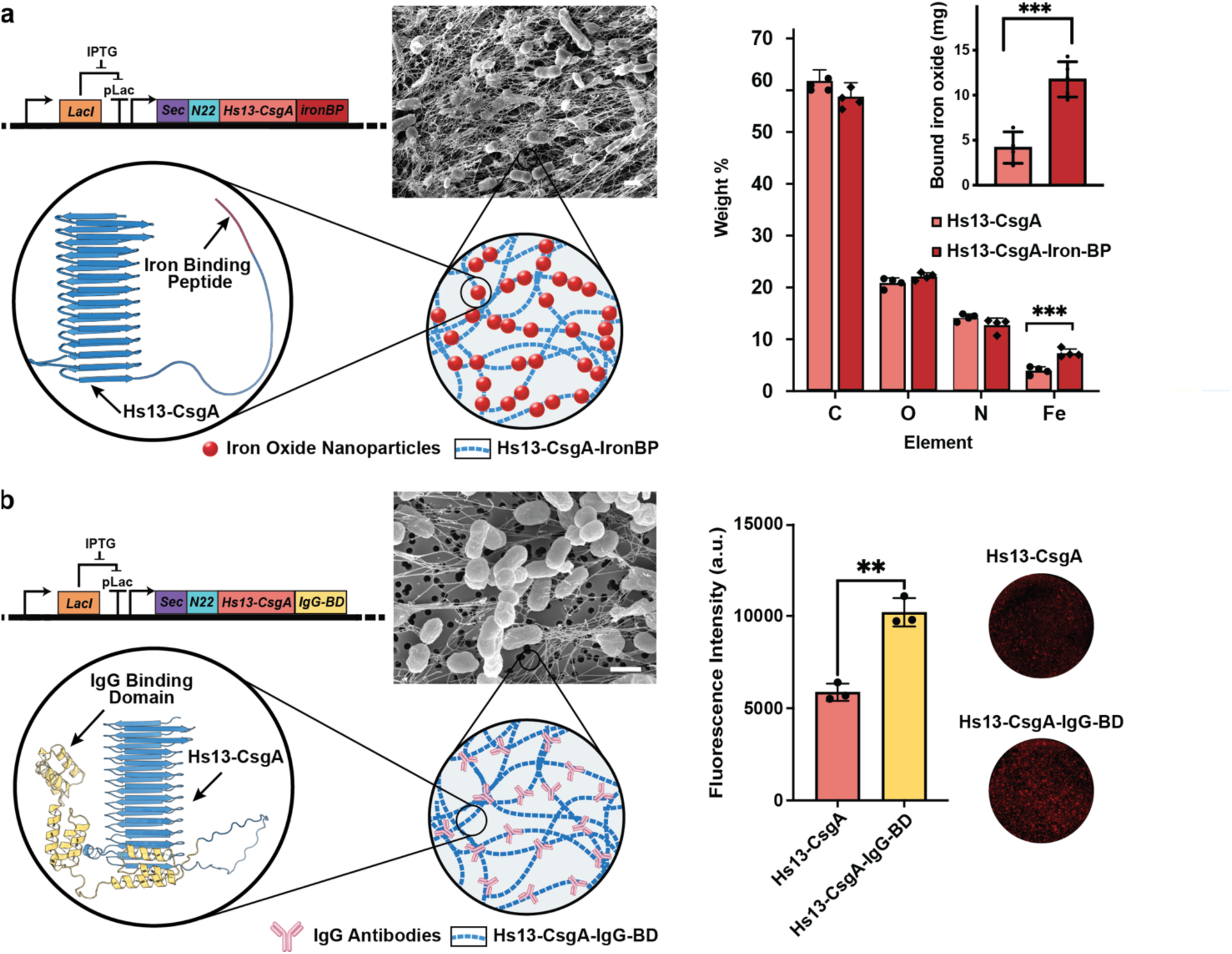
Programming the biological functionalities of *de novo* β-solenoid proteins. **a** Genetic design of *E. coli* programmed to produce β-solenoid protein nanofibers displaying iron oxide binding peptide (Hs13-CsgA-IronBP) along with FESEM image of the protein nanofibers self-assembled in the extracellular milieu. Weight analysis and energy dispersive X-ray (EDAX) analysis show the enhanced binding of iron oxide nanoparticles to engineered nanofibers, Hs13-CsgA-IronBP. Scale bar 1 μm. Biological replicates n = 4. Data represented as mean ± standard deviation. ***p≤0.0001 two-way ANOVA for EDAX analysis and ***p≤0.0008 t-test followed by Welch’s test for weight analysis. **b** Genetic design of *E. coli* cells to produce β-solenoid protein nanofibers displaying IgG antibody binding domain (Hs13-CsgA-IgG-BD) along with FESEM image of the protein nanofibers self-assembled in the extracellular milieu. Fluorescent IgG antibodies binding assay shows higher binding for Hs13-CsgA-IgG-BD than wildtype Hs13-CsgA nanofibers. Scale bar 1 μm. Biological replicates n = 3. Data represented as mean ± standard deviation. **p≤0.0024 t-test followed by Welch’s test. The structures of *de novo* β-solenoid proteins, Hs13-CsgA-IronBP and Hs13-CsgA-IgG-BD were predicted using AlphaFold2.

## Discussion

There are only a few examples of functional amyloids, and therefore, it was initially surprising for us to find that 33,564 proteins listed in the NCBI database are homologs of CsgA. Interestingly, recent work has reported that 22% (43279) of the genomes out of the 201210 bacterial genomes searched contained one or more predicted CsgA sequences with one or more curli repeat signatures.^32, 41^ Thus, our protein mining results corroborate closely with the previously reported genome mining results and, on the other hand, emphasize the enormous scale to which CsgA-like proteins are prevalent in the natural world.^41^ Moreover, the curli system is not exclusively limited to *E. coli*, and ∼33% of identified CsgA homologs are secreted by bacteria phylogenetically spread across at least four major bacterial phyla, each with its own distinct curli system. This evolutionary diversity and abundance of functional proteins highlight the untapped potential that could be explored and exploited for advanced autogenic ELMs.

The rapid advancement of AI-driven structure prediction tools such as AlphaFold2 has revolutionized our ability to explore the structural properties of diverse proteins. These tools have significantly accelerated the design, screening, and development of novel functional protein-based materials. As cryo-TEM (transmission electron microscopy) and single crystal X-ray diffraction studies of CsgA and its homologs have proven extremely difficult, structure prediction tools like AlphaFold2 are beneficial in getting critical insights. MD simulations provide valuable insights, demonstrating the critical role of KRs in stabilizing β-solenoid structure. However, the evolutionary diversity observed in *de novo* analogs indicates that even non-conserved KR_1_ and KR_7_ residues lead to stable structures. Thus, it opens a broad landscape for tinkering and rationally designing *de novo* functional protein materials.

In summary, we have demonstrated that *E. coli* curli machinery could be meticulously utilized to produce *de novo* autogenic ELMs by employing β-solenoid protein building blocks of non-model extremophilic bacteria. Despite having distinct amino acid compositions and molecular weights (up to 9 times of native CsgA), their biosynthesis, secretion, and extracellular self-assembly into stable β-solenoid protein nanofibers highlight the robustness and versatility of the SECRETE platform to biomanufacture *de novo* autogenic ELMs. Moreover, the variations in the yields of hydrogels (375 to 1938 mg L^−1^) and storage modulus (335 to 959 Pa) might be influenced by the physicochemical properties of β-solenoid building blocks, such as the number of β-sheet repeats, hydrophobicity, charge, and amino acid compositions, which needs further investigation. The fabrication of hydrogels, films, and well-defined 3D architectures with programmable physical, chemical, and biological functionalities will open Pandora’s box to rationally design sophisticated *de novo* autogenic ELMs for biomedical and environmental applications.

## Contributions

A.D.T. and A.M.B. developed the initial concept. A.D.T. streamlined the central concept and methodology. A.D.T. and H.H. designed experiments. H.H. performed protein mining, AlphaFold2 structure prediction, and engineering of *E. coli* to produce β-solenoid protein nanofibers, hydrogels, and their characterization. S.S. performed all molecular dynamics simulations and all analyses. E.C. performed WAXS analysis of β-solenoid proteins. A.M.B. performed SEM analysis. R.M. supervised E.C.; S.D. supervised S.S.; A.D.T. supervised H.H. The manuscript was drafted by H.H., A.M.B., and A.D.T. with input from all authors. All authors discussed the results and commented on the manuscript.

## Acknowledgment

Work was partly performed at the Advanced Research Computing Center, Nanoscale Characterization and Fabrication Laboratory (NCFL), and Institute for Critical Technology and Applied Science (ICTAS) at Virginia Tech. This work was supported by GlycoMIP, a National Science Foundation Materials Innovation Platform funded through Cooperative Agreement DMR-1933525.

## Materials and Methods

### Cell strains and plasmids

All experiments were conducted using PQN4, *E. coli* strain derived from LSR10 (MC4100, ΔcsgA, λ(DE3), CamR) with the deletion of curli operon (ΔcsgBACEFG).^42^ The genes encoding CsgA and its 5 homologs and the two engineered domains were synthesized by Twist BioScience and cloned into the pET21d vector using overlap extension and isothermal Gibson Assembly (New England Biolabs). The whole plasmid sequences were confirmed by Plasmidsaurus. These plasmids also included genes encoding proteins essential for curli biosynthesis, such as csgC, csgE, csgF, and csgG. Five different β-solenoid proteins, namely, Hs13-CsgA, Am18-CsgA, Bc36-CsgA, El43-CsgA, and Er46-CsgA were inserted in place of the *E. coli* wild-type CsgA after the SEC (N-terminal signal sequence) and the N22 (N-terminal curli-specific targeting sequence) to facilitate the secretion of CsgA variants into the extracellular space. Additionally, two engineered β-solenoid protein variants were created, namely, Hs13-CsgA-IronBP and Hs13-CsgA-IgG-BD by fusing iron-binding peptide (IronBP) and antibody IgG binding domain (IgG-BD) to the C-terminus of Hs13-CsgA via a 36-amino-acid flexible linker. All gene sequences, protein sequences, and primers used in this study are fully provided in **Supplementary Fig. 17** and **Supplementary Table 4**.

### Protein sequence mining and AlphaFold2 modeling of CsgA and its homologs

We utilized the protein structure database of the National Center for Biotechnology Information (NCBI) and employed the search query - Major [Organism] OR major [All Fields] AND curlin [All Fields] - to identify CsgA homologs. This search yielded 33,564 entries, annotated as having the curlin subunit CsgA, either through computational analysis or by identifying sequences using a regular expression based on the minimalistic curli repeat (X_6_QXGX_2_NX_10_) described by Chapman et al.^43^ The lengths of the resulting protein sequences varied from 141 to 1390 amino acids. From these 33,564 entries, we randomly selected 50 sequences, and to predict their 3D structure, we employed AlphaFold2 (version 2.1.1)^31^ on the Tinkercliffs HPC cluster at the Advanced Research Computing Center, Virginia Tech, utilizing NVIDIA A100-80G architecture. The source code for AlphaFold2 is available at https://github.com/deepmind/alphafold. Structural images were generated using PyMOL (version 1.16).^44^ Batch protein structure predictions were performed, running for six recycles and generating five models per sequence. The models were ranked using the pLDDT score, with the highest-ranked model being used for subsequent analyses. Out of these 50 sequences, 5 sequences were selected, having lengths across the whole spectrum of 141 to 1390 amino acids. Genes of these 5 sequences were synthesized to validate their production in the *E. coli* PQN4 strain experimentally. These variants were named based on the first letter of the species, followed by the number of cross-β amyloid repeats and CsgA - to indicate that it is a homolog. For example, the CsgA homolog of *Halomonas saliphila* with 13 cross-β amyloid repeats was named Hs13-CsgA. Similarly, 18, 36, 43, and 46 cross-β amyloid repeats of CsgA homologs of *Alteromonas macleodii*, *Blastomonas sp*. CACIA14H2, *Erythrobacter longus*, and *Ensifer sp. Root31* were labelled as Am18-CsgA, Bc36-CsgA, El43-CsgA and Er46-CsgA, respectively. In addition, two engineered variants, Hs13-CsgA-IronBP and Hs13-CsgA-IgG-BD, were also predicted in a similar fashion.

### Molecular dynamics (MD) simulation

All-atom MD simulations of the six β-solenoid proteins – CsgA, Hs13-CsgA, Am18-CsgA, Bc36-CsgA, El43-CsgA, and Er46-CsgA in explicit water were conducted using the NAMD-2.14 software.^45^ The initial structures were obtained from the AlphaFold2 server using the experimental FASTA sequences.^31^ These β-solenoid proteins were solvated using TIP3P water in a cubic box with a minimum of 15 Å padding in each direction. The solvated β-solenoid proteins were neutralized by adding sodium ions to balance the net charges of these proteins (**Supplementary Figure 7**). The atomistic MD simulations were performed in 2 fs timestep in the NPT ensemble with the periodic boundary conditions applied in all directions. The Charmm-36m (July 22) force field (FF) was used to model the bonded and non-bonded interactions.^46^ The standard 12-6 Lennard-Jones potential was applied with a 12 Å cutoff, and the Particle Mesh Ewald (PME) method was used to calculate the long-range electrostatic interactions.^47^ The Langevin thermostat and Parrinello-Rahman barostat were implemented to maintain a temperature at 300 K and pressure at 1.01325 bar, respectively. This simulation setup maintained the system stability while accommodating fluctuations in density. Energy minimization of 50,000 steps was followed by 150 ns MD simulations, in which the last 50 ns were treated as a production run. The trajectory data was stored in 1 ps intervals during these MD simulations. All the analyses were carried out using the last 50 ns production run except for the RMSDs, which were calculated for the entire 150 ns trajectory to compare and estimate the structural changes from the initial AlphaFold2 generated structures.

### Analysis method of MD simulation

The RMSD and RMSF of the amyloids were calculated using the VMD software^48^ package, and the MD Analysis package was used to calculate the hydrogen bond autocorrelations and Ramachandran plots. The hydrogen bond criteria used were a donor-hydrogen distance cutoff of 1.2 Å, a donor-acceptor distance cutoff of 3.0 Å, and a donor-hydrogen-acceptor angle cutoff of 150°.^36^ We developed an automated tool utilizing the NAMD executable to calculate the residue-residue interaction energies for the large β-solenoid protein variants (https://github.com/Deshmukh-Group/ResidueResidue_Interaction_Energies_NAMD). These energies were calculated based on the non-bonded interactions, including electrostatic and van der Waals contributions. Water and ions were removed from the system to calculate the energies. The last 50 ns trajectory is analyzed with a 0.1 ns stride. A percent cutoff of 60% and 12 Å filtering distance were applied, meaning that only pairs of residues whose center-of-mass come closer than 12 Å in at least 60 percent of trajectory frames were included in the non-bonded energy calculations. We verified our energy data with the gRINN software package.^49^

### Microbial production of β-solenoid protein nanofibers

All plasmids (CsgA, Hs13-CsgA, Am18-CsgA, Bc36-CsgA, El43-CsgA, Er46-CsgA, Hs13-CsgA-IronBP, and Hs13-CsgA-IgG-BD) were transformed into the *E. coli* strain PQN4. The transformed cells were streaked onto lysogeny broth (LB) agar plates containing 100 µg ml^−1^ carbenicillin and 0.5% glucose (w v^−1^) for catabolite repression of T7RNAP and incubated overnight at 37 °C. A single colony from each plate was picked and separately cultured at 37 °C in 5 ml of LB media with 100 µg ml^−1^ carbenicillin and 2% glucose (w v^−1^). These overnight cultures (PQN4_CsgA, PQN4_Hs13-CsgA, PQN4_Am18-CsgA, PQN4_Bc36-CsgA, PQN4_El43-CsgA, PQN4_Er46-CsgA, PQN4_Hs13-CsgA-IronBP, and PQN4_Hs13-CsgA-IgG-BD) were then transferred to fresh 500 ml LB media containing 100 µg ml^−1^ carbenicillin. The cultures were incubated in shaking incubators (225 rpm, 37 °C) for 48 hours to achieve expression of β-solenoid proteins and assembly into functional amyloid nanofibers. As a negative control, PQN4 was transformed with a sham (all curli genes, including csgA were absent), and pET21d plasmid was used.

### Quantitative Congo red dye binding assay

One milliliter of bacterial culture (48h, 500 ml) was centrifuged at 6000 rpm for 10 minutes. The resulting cell pellet was resuspended in a 0.025 mM solution of Congo red in phosphate-buffered saline (PBS) and incubated for 10 minutes. The cells were then pelleted again at 14,000 rpm for 10 minutes, and the absorbance of the supernatant (200 µl) at 490 nm was measured using a microplate reader. This absorbance value was subtracted from 0.025 mM Congo red in PBS, which was subsequently normalized by the OD600 of the original bacterial culture to quantify the production of functional amyloid nanofibers. The assay was performed in four biological replicates.

### Congo Red birefringence

Functional amyloid nanofibers were stained with Congo red and evaluated for birefringence.^50^ One milliliter of bacterial culture was centrifuged at 6000 rpm for 10 minutes. The resulting pellet was resuspended in 10 μl of 500 µM Congo red solution in 80% ethanol and incubated for 60 minutes at room temperature. XploRA™ PLUS Confocal Raman Microscope, equipped with an inverted camera and crossed polarizing light filters, was utilized to visualize and assess birefringence.

### Preparation of β-solenoid protein hydrogels

*E. coli* PQN4 strain with the desired plasmid was cultured in 500 ml of LB media in a shaking incubator at 37 °C, 225 rpm for 48 hours to produce the β-solenoid protein nanofibers. The resulting 500 ml culture was treated with urea to a final concentration of 0.8 M and kept at 4 °C for 1 h. The treated cultures were then concentrated by vacuum filtration through a 90-mm diameter polycarbonate membrane with 10-μm pores (EMD Millipore). The resulting concentrated biofilm (nanofibers) was washed thrice with 50 ml of sterile deionized (DI) water on the filter membrane. The biofilm was subsequently incubated with 100 ml of 8M urea solution in water for 5 minutes, followed by vacuum filtration and washing with 200 ml of DI water to remove the bacterial debris lysate. The resulting biomass on the filter membrane was then treated with 50 ml of 5% (w v-1) SDS (sodium dodecyl sulfate) solution (gelator/plasticizer) in water for 5 minutes, followed by vacuum filtration and an additional wash with 500 ml of DI water. The β-solenoid protein hydrogel formed on the filter membrane was then collected and stored at 4 °C.

### Fabrication of aquaplastic from β-solenoid protein nanofibers

To fabricate 2D aquaplastic films, β-solenoid protein nanofibers based hydrogels were cast onto a flat, flexible substrate such as plastic wrap or aluminum foil. These were left to dry overnight under ambient conditions to form aquaplastic films of lateral dimensions 1 cm * 1 cm. Once thoroughly dried, the aquaplastic film was gently peeled off the substrate and stored at room temperature.

### Field-emission scanning electron microscopy sample preparation and imaging

100 µl of the *E. coli* cultures or 10 mg of hydrogel samples were filtered/spread onto polycarbonate membranes with a 0.22 μm pore size under vacuum and then placed in a fixative solution containing 1 ml of 4% glutaraldehyde and 1 ml of 4% paraformaldehyde buffer for 2 hours at room temperature. After fixation, the membranes were gently rinsed with water and subjected to a series of 200-proof ethanol washes with increasing concentrations (25%, 50%, 75%, 100%, and 100% v v-1), each for 15 minutes. The samples were then processed using a critical point dryer. The dried membranes were mounted on Scanning Electron Microscopy sample holders with carbon adhesives and coated with a 10 to 20 nm layer of Pt/Pd. Images were captured using a JEOL IT500 SEM equipped with a field-emission gun operating at 5-10 kV.

### Wide-angle X-ray scattering

Wide-angle X-ray scattering (WAXS) experiments on the aquaplastics were performed using a Xenocs Xeuss 3.0 SAXS/WAXS equipped with a GeniX 3D Cu HFVLF microfocus X-ray source utilizing Cu Kα radiation (λ = 0.154 nm). The sample-to-detector distance was 42 mm, and the q-range was calibrated using a Lanthanum hexaboride standard. Two-dimensional scattering patterns were obtained using a Dectris EIGER 4M detector with an exposure time of 30 minutes. Data reduction was performed using XSACT software provided by Xenocs.

### Rheology studies of the hydrogels

The viscoelastic properties of the β-solenoid protein hydrogels were assessed using a µ-volume sample holder with the ElastoSens™ Bio (Rheolution, Montreal, Canada). 250 µg gel sample was cast into the sample holder. The storage modulus (G′, Pa) of each sample was measured to evaluate their viscoelastic characteristics. Each test was performed in technical duplicates for three biological replicates, resulting in a total of six readings per time point. G′ was recorded every 5 seconds over a 1-minute period.

### 3D printing of β-solenoid protein hydrogels (Microbial Ink)

The bioinks were transferred into a 10-ml Luer-Lok™ syringe and centrifuged at 112 × g for 2 minutes to remove any air bubbles. Bioprinting was conducted at room temperature using a Hyrel HYDRA 21 Bioprinter (Hyrel3D, Norcross, GA, USA) equipped with a 10-ml syringe dispensing head (SDS-10) for gel printing. 22G and 20G needles with a premade 0.25-inch blunt end were utilized. The printing speed of 2 mm s^−1^ and the blunt needle diameters of 0.41 mm (22G) and 0.60 mm (20G) were selected after optimization. This process involved assessing various printing speeds and needle sizes with internal diameters ranging from 0.26 to 0.60 mm to achieve optimal pattern fidelity, reproducibility, and consistency across different polymer concentrations and viscosities of the amyloid hydrogel inks. Initial bioprinting trials were performed at feed rates ranging from 2 to 10 mm s^−1^. The 3D STL files were prepared using the slic3r engine integrated with the Hyrel HYDRA 21 Bioprinter software.

### Optical Images

Optical images were obtained using a Canon EOS rebel T7 DSLR Camera.

### Iron oxide binding analysis by genetically engineered β-solenoid protein

The plasmid pET21d-Hs13-CsgA-IronBP and pET21d-Hs13-CsgA were separately transformed into PQN4 cells to produce engineered and non-engineered functional amyloid nanofibers. 50 ml of culture with expressed nanofibers was spun down, and the supernatant was discarded. This cell pellet was added with 5 mg of iron (III) oxide (<50 nm, Sigma-Aldrich) in 50 ml of water. Cell pellets with nanofibers were incubated at ambient temperature with light agitation for 2 hours to facilitate binding of iron oxide nanoparticles to nanofibers. Next, the entire mixture was deposited on a 47-mm diameter polycarbonate membrane with 10-μm pores (EMD Millipore) and washed with water several times to remove unbound iron oxide nanoparticles. The weight of iron oxide nanoparticles bound to Hs13-CsgA-IronBP was determined and compared to control samples with non-engineered nanofibers of Hs13-CsgA. Additionally, the engineered and non-engineered nanofibers with bound iron oxide nanoparticles deposited on the filter membrane were tested by energy-dispersive X-ray analysis (EDAX) to obtain elemental composition.

### Antibody binding analysis of genetically engineered β-solenoid protein

The plasmid pET21d-Hs13-CsgA-IgG-BD was transformed into PQN4 cells to express the engineered functional amyloids containing IgG binding protein fusion. The expressed engineered nanofibers were deposited on the filter membrane (dry weight of nanofibers 1.875 ± 0.62 mg) and incubated with a solution of antibody Goat anti-Rabbit IgG with Alexa Fluor™ 594 (Thermo Fisher Scientific) for 2 hours. Next, filter membranes with deposited, engineered (Hs13-CsgA-IgG-BD) and non-engineered nanofibers (control, Hs13-CsgA) were washed with water several times to remove unbound antibodies. The bound antibodies were detected by fluorescence intensity of Alexa Fluor 594 using ChemiDoc Imaging System (BIO-RAD). Images were analyzed by utilizing ImageJ software.

### Statistics and reproducibility

All experiments presented in this article were repeated at least three times (n=3) on biological replicates or distinct samples, and they are clearly stated in the figure captions and relevant method sections. All data are presented as the mean and standard deviation. All plotting and statistical analysis (ordinary one-way ANOVA or Welch t-test) were performed using GraphPad Prism 10 software.

## Data availability

The article and its supplementary information contain all relevant data supporting this study’s findings. Source data are provided in this paper.

