## Supplementary Information for "*De novo* Autogenic Engineered Living Functional Materials"

**Supplementary Figure 1. Genetically modified CsgA structure prediction using AlphaFold2.** The protein structure of CsgA, with C-terminal fusion of Trefoil Factor 2 (TFF2) predicted by AlphaFold2, shows the β-solenoid fold of CsgA.

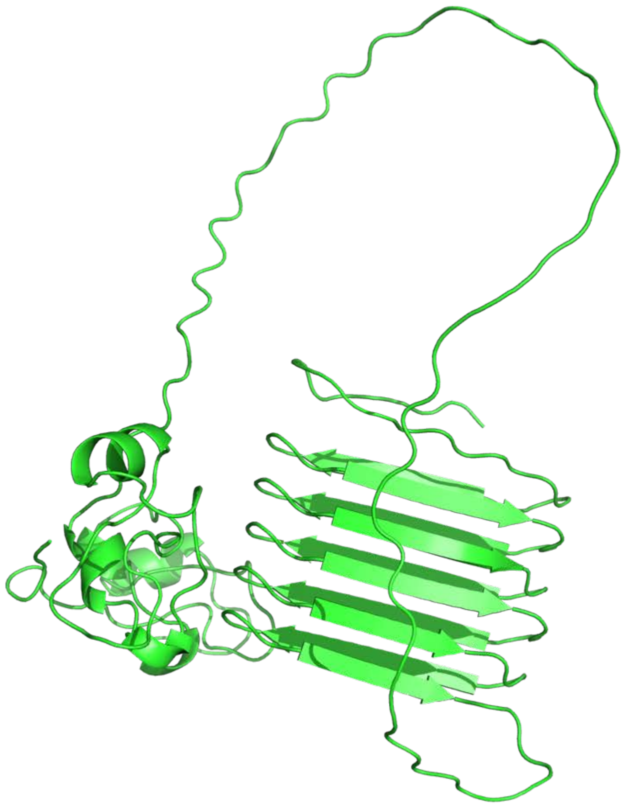

**Supplementary Figure 2**. **AlphaFold2 structure prediction of β-solenoid proteins.** Predicted structure of 50 random sequences obtained from protein mining. The prediction shows β-solenoid fold with cross-β repeat units ranging from 5 to 46.

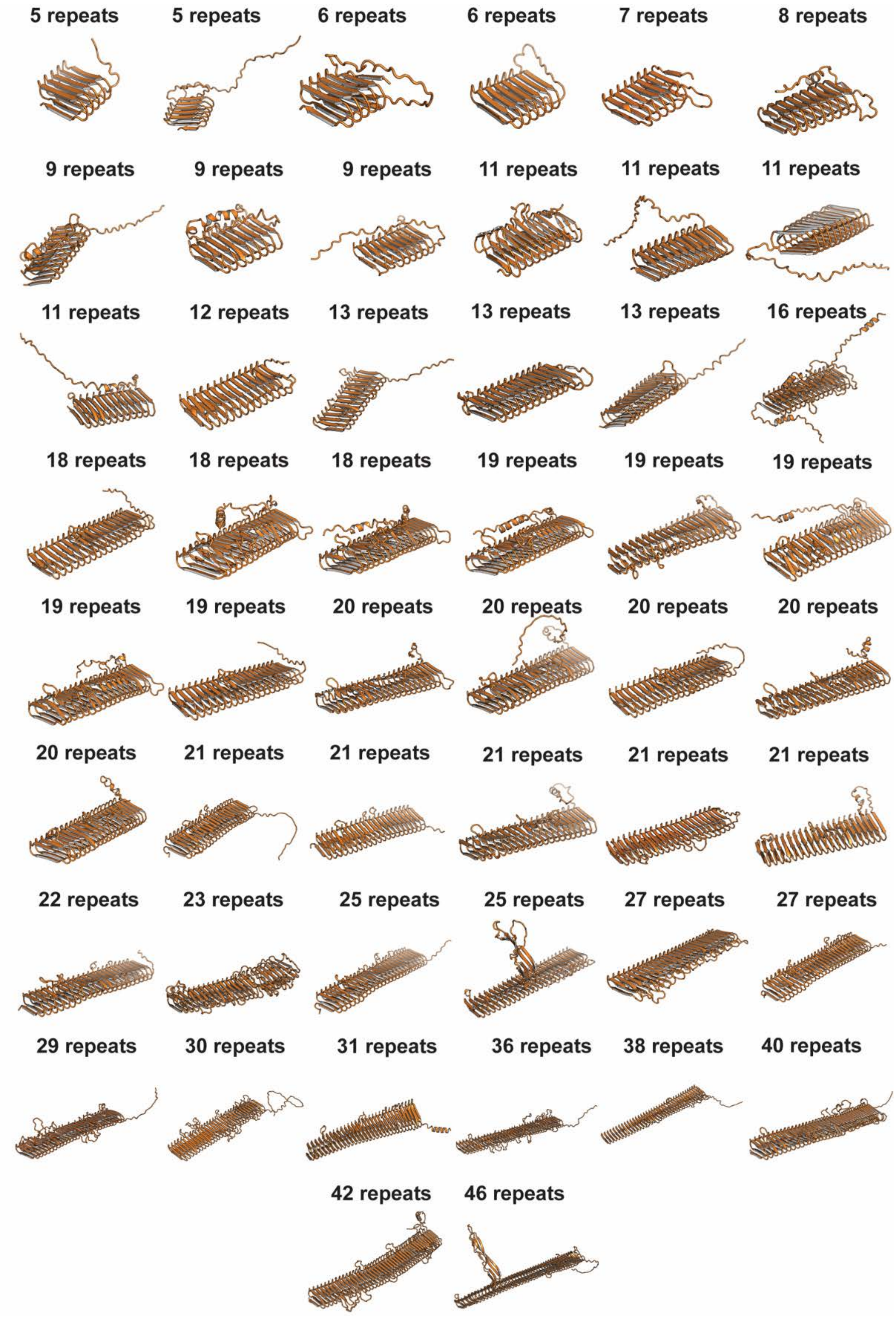

**Supplementary Figure 3. Sequence alignment of β-solenoid proteins used in this study.** The protein sequence alignment of CsgA, Hs13-CsgA, Am18-CsgA, Bc36-CsgA, El43-CsgA and Er46-CsgA, show β-strands (β1 and β2), loops (intra-repeat and inter-repeat) and outlier region (Σ).

| **CsgA** | β1 strand | | | | | | | Intra-Repeat Loops | | | | β2 strand | | | | | | | Inter-Repeat Loops | | | | | |
|---|---|---|---|---|---|---|---|---|---|---|---|---|---|---|---|---|---|---|---|---|---|---|---|---|
| | S | Ω | Ψ | Ω | Ψ | Ω | Q | x | G | x | G | N | Ω | Ψ | Ω | Ψ | Ω | Q | x | x | x | x | | |
| N-terminal motif | | | G | V | V | P | Q | Y | G | G | G | G | N | H | G | G | G | G | N | N | S | G | P | N |
| Repeat Unit 1 | S | E | L | N | I | Y | Q | Y | G | G | G | N | S | A | L | A | L | Q | T | D | A | R | N | |
| Repeat Unit 2 | S | D | L | T | I | T | Q | H | G | G | G | N | G | A | D | V | G | Q | G | S | D | D | | |
| Repeat Unit 3 | S | S | I | D | L | T | Q | R | G | F | G | N | S | A | T | L | D | Q | W | N | G | K | N | |
| Repeat Unit 4 | S | E | M | T | V | K | Q | F | G | G | G | N | G | A | A | V | D | Q | T | A | S | N | | |
| Repeat Unit 5 | S | S | V | N | V | T | Q | V | G | F | G | N | N | A | T | A | H | Q | Y | | | | | |

| **Hs13-CsgA** | β1 strand | | | | | | | Intra-Repeat Loops | | | | | β2 strand | | | | | | | Inter-Repeat Loops | | | | |
|---|---|---|---|---|---|---|---|---|---|---|---|---|---|---|---|---|---|---|---|---|---|---|---|---|
| | N | Ω | Ψ | Ω | Ψ | Ω | Q | x | G | x | x | Σ | N | Ω | Ψ | Ω | Ψ | Ω | Q | x | G | x | x | Σ |
| N-terminal motif | M | K | L | K | M | T | V | M | A | A | A | | I | A | V | S | S | T | A | L | A | N | E | |
| Repeat Unit 1 | N | E | S | N | I | A | Q | T | G | F | G | | N | D | V | S | V | T | Q | L | T | Q | G | WGN |
| Repeat Unit 2 | N | N | S | D | V | T | Q | V | G | V | N | | N | G | A | T | V | L | Q | Q | G | W | R | GR |
| Repeat Unit 3 | N | V | S | D | I | D | Q | E | G | M | G | | N | N | A | D | A | L | Q | H | G | T | K | |
| Repeat Unit 4 | N | S | F | D | L | T | Q | S | G | M | F | | N | N | S | D | S | Q | Q | L | G | T | | |
| Repeat Unit 5 | N | A | L | R | V | E | Q | E | G | V | K | | N | N | A | T | T | Y | Q | S | G | W | R | |
| Repeat Unit 6 | N | Y | A | D | I | S | Q | S | G | S | F | | N | N | S | D | V | D | Q | E | G | Y | L | |
| Repeat Unit 7 | N | V | A | Q | V | N | Q | D | G | T | K | | L | S | S | D | I | T | Q | S | G | D | Y | |
| Repeat Unit 8 | N | F | A | D | V | D | Q | S | G | S | F | | N | D | S | S | I | D | Q | N | G | T | Q | |
| Repeat Unit 9 | N | K | A | E | V | L | Q | S | G | L | K | | N | T | S | D | I | H | Q | D | G | K | L | |
| Repeat Unit 10 | N | I | A | M | V | D | Q | S | G | V | N | | N | D | A | S | I | V | Q | D | G | R | Y | |
| Repeat Unit 11 | H | Q | S | E | V | T | Q | S | G | Y | N | | D | V | A | Y | S | S | Q | S | G | Y | E | |
| Repeat Unit 12 | H | Y | S | S | I | T | Q | S | A | L | P | GFFGGIASIAS | N | S | A | T | H | V | Q | M | G | V | N | |
| Repeat Unit 13 | N | A | A | I | T | T | Q | V | G | G | G | | N | T | A | H | V | Y | Q | H | | | | |

| Am18-CsgA | β1 strand | | | | | | | Intra-Repeat Loops | | | | | β2 strand | | | | | | | Inter-Repeat Loops | | | | | | | | | |
|---|---|---|---|---|---|---|---|---|---|---|---|---|---|---|---|---|---|---|---|---|---|---|---|---|---|---|---|---|---|
| | N | Ω | Ψ | Ω | Ψ | Ω | X | x | G | x | x | Σ | N | Ω | Ψ | Ω | Ψ | Ω | Q | x | G | x | x | Σ | | | | | |
| N-terminal motif | M | K | M | K | K | T | K | I | A | S | A | | I | I | I | V | A | A | T | G | W | A | S | A | Q | E | I | P | A |
| N-terminal motif | S | K | S | V | D | F | N | A | S | A | D | | F | V | A | A | A | E | G | | | | | | | | | | |
| Repeat Unit 1 | N | E | I | T | L | T | Q | T | A | P | E | GNEVG | N | E | S | V | L | S | Q | E | G | D | L | | | | | | |
| Repeat Unit 2 | N | A | I | V | V | E | V | T | G | D | A | | N | E | V | L | A | S | Q | M | Q | S | G | | | | | | |
| Repeat Unit 3 | N | T | F | F | A | S | V | T | G | N | G | | N | F | L | E | S | E | Q | D | S | L | D | | | | | | |
| Repeat Unit 4 | S | T | A | E | F | V | V | E | G | D | D | | N | L | V | S | L | V | Q | L | G | D | G | GPFGFIF | | | | | |
| Repeat Unit 5 | S | R | S | I | N | S | I | A | G | S | G | | N | T | L | S | V | Y | Q | G | D | G | G | | | | | | |
| Repeat Unit 6 | N | W | A | N | N | A | I | A | G | N | D | | N | T | V | F | V | D | Q | S | G | D | W | | | | | | |
| Repeat Unit 7 | H | E | S | Y | V | R | E | L | S | G | D | A | N | L | I | D | V | I | Q | D | G | F | Y | | | | | | |
| Repeat Unit 8 | N | I | S | D | L | T | V | V | G | S | D | | N | E | V | E | V | D | Q | D | G | D | E | | | | | | |
| Repeat Unit 9 | N | L | I | T | W | N | M | V | G | D | G | | N | V | L | E | F | E | Q | D | G | D | G | | | | | | |
| Repeat Unit 10 | N | E | I | T | T | G | V | F | E | G | S | D | N | E | V | N | I | D | Q | I | G | D | V | | | | | | |
| Repeat Unit 11 | N | L | A | T | V | E | T | I | G | G | G | V | N | T | F | E | I | N | Q | V | G | E | G | | | | | | |
| Repeat Unit 12 | N | V | A | Y | A | G | V | I | G | L | F | | N | E | F | D | L | T | Q | V | G | D | A | | | | | | |
| Repeat Unit 13 | N | E | I | S | T | V | N | F | D | G | W | F | N | T | V | E | I | E | Q | D | G | D | A | | | | | | |
| Repeat Unit 14 | N | L | A | L | A | Q | A | G | I | G | D | AYVADA | N | L | I | E | M | S | Q | E | G | N | E | | | | | | |
| Repeat Unit 15 | N | I | A | S | V | E | L | A | R | D | I | TSGS | N | E | I | M | V | A | Q | T | G | E | L | | | | | | |
| Repeat Unit 16 | N | L | L | D | L | L | V | N | G | N | D | | N | V | I | S | V | M | Q | E | G | A | G | | | | | | |
| Repeat Unit 17 | N | W | V | T | D | D | M | G | G | Q | F | VISGDM | N | T | F | E | V | T | Q | M | G | N | D | | | | | | |
| Repeat Unit 18 | N | L | V | T | G | S | I | T | G | N | G | | G | T | V | S | V | T | Q | V | G | D | Y | | | | | | |
| Half Repeat Unit | N | V | A | T | V | V | Q | M | | | | | | | | | | | | | | | | | | | | | |

| Bc36-CsgA | β1 strand | | | | | | | Intra-Repeat Loops | | | | | β2 strand | | | | | | | Inter-Repeat Loops | | | | |
|---|---|---|---|---|---|---|---|---|---|---|---|---|---|---|---|---|---|---|---|---|---|---|---|---|
| | N | Ω | Ψ | Ω | Ψ | Ω | Q | x | G | x | x | Σ | X | Ω | Ψ | Ω | Ψ | Ω | Q | x | G | x | x | Σ |
| N-terminal motif | M | K | T | S | I | Y | T | S | V | S | A | | I | A | I | M | I | G | M | P | A | V | A | QT |
| Repeat Unit 1 | N | T | S | T | V | D | Q | T | G | A | A | | A | A | A | T | V | N | Q | T | G | S | N | |
| Repeat Unit 2 | N | T | S | D | I | D | Q | N | G | N | G | IAYTTGPNTGRT | V | V | A | D | V | E | Q | T | G | S | N | |
| Repeat Unit 3 | G | F | S | Q | V | T | Q | Q | G | G | R | | S | S | A | T | V | D | Q | G | G | T | G | |
| Repeat Unit 4 | M | R | S | T | I | T | Q | G | N | A | S | TDIG | N | T | A | T | V | I | Q | N | G | T | G | GGVG |
| Repeat Unit 5 | T | G | S | T | I | T | Q | T | G | G | T | | G | Q | A | Y | V | N | Q | G | A | A | T | NG |
| Repeat Unit 6 | A | I | S | T | I | T | Q | T | G | N | N | | Q | R | A | S | V | F | Q | T | S | G | A | |
| Repeat Unit 7 | A | D | S | S | V | S | Q | S | G | G | A | | A | N | V | F | V | S | Q | D | G | S | . | |
| Repeat Unit 8 | S | E | S | D | I | T | Q | T | G | G | N | | S | E | A | S | V | R | Q | I | G | D | G | |
| Repeat Unit 9 | N | T | S | L | I | E | Q | T | G | V | N | GDVGDP | D | N | N | L | A | N | Q | D | T | N | I | GVSQTGDN |
| Repeat Unit 10 | N | S | S | T | V | R | Q | T | G | N | D | | Q | V | A | D | V | I | Q | T | G | D | F | |
| Repeat Unit 11 | N | T | A | S | I | T | Q | D | G | N | F | | A | D | A | S | V | T | Q | T | G | N | N | N |
| Repeat Unit 12 | T | G | T | V | I | R | Q | D | G | D | G | SGDNDPASIPVIPTTN | A | T | A | D | I | S | Q | T | G | D | F | |
| Repeat Unit 13 | N | E | A | S | I | S | Q | A | A | V | P | | V | E | A | T | I | T | Q | L | G | D | S | |
| Repeat Unit 14 | N | D | S | S | I | V | Q | S | S | T | A | SG | A | K | A | T | N | L | Q | T | S | D | N | |
| Repeat Unit 15 | N | L | S | T | I | T | Q | D | S | D | . | | A | E | A | S | V | T | Q | G | G | S | Q | FNPGPFAGRDN |
| Repeat Unit 16 | N | V | S | T | V | A | Q | G | T | G | A | AG | S | L | A | T | V | N | Q | D | G | V | L | |
| Repeat Unit 17 | N | L | S | D | I | N | Q | N | A | A | N | | S | T | A | S | V | T | Q | T | N | I | F | |
| Repeat Unit 18 | N | T | S | S | V | I | Q | S | G | S | G | G | A | T | A | D | V | T | Q | G | G | G | Y | GD |
| Repeat Unit 19 | N | V | S | N | I | T | Q | T | T | N | . | | A | S | A | T | V | V | Q | T | G | Q | P | SVGPDFT |
| Repeat Unit 20 | N | I | S | D | I | V | Q | N | G | T | G | | G | I | A | S | V | T | Q | N | G | S | N | |
| Repeat Unit 21 | N | E | S | D | V | A | Q | G | G | T | D | | G | D | A | T | V | D | Q | D | G | T | F | |
| Repeat Unit 22 | L | R | S | T | I | T | Q | T | G | G | E | | A | T | A | L | V | T | Q | T | G | N | A | |
| Repeat Unit 23 | N | V | S | T | I | S | Q | S | A | A | S | AG | A | D | A | T | V | T | Q | S | G | G | E | |
| Repeat Unit 24 | N | R | S | D | V | N | Q | T | G | A | . | | A | K | A | V | V | T | Q | A | G | T | E | VGYNIAPPN |
| Repeat Unit 25 | N | D | S | F | V | V | Q | S | A | D | G | | A | D | A | N | V | S | Q | T | G | E | L | |
| Repeat Unit 26 | N | L | S | N | V | N | Q | S | G | G | A | | S | E | A | D | V | T | Q | T | G | V | Y | |
| Repeat Unit 27 | N | R | S | T | V | T | Q | T | V | G | G | | A | K | A | T | V | T | Q | S | G | D | P | ANGGPGVGQGD |
| Repeat Unit 28 | N | L | S | I | V | S | Q | S | G | A | . | | S | T | A | T | V | T | Q | T | A | N | V | PTNGFPS |
| Repeat Unit 29 | N | D | S | F | V | S | Q | T | G | D | D | | S | D | A | T | V | D | Q | T | G | N | D | |
| Repeat Unit 30 | N | A | S | D | V | F | Q | G | S | D | . | | A | I | A | D | V | T | Q | D | G | T | N | |
| Repeat Unit 31 | N | T | S | T | V | T | Q | S | A | A | . | | T | S | A | F | V | E | Q | I | G | A | R | |
| Repeat Unit 32 | N | T | S | T | V | T | Q | S | G | G | S | TVAP | Y | A | A | Y | V | Q | Q | L | G | N | D | |
| Repeat Unit 33 | G | L | S | V | V | E | Q | D | G | A | N | | N | E | A | T | L | T | Q | S | A | G | S | SF |
| Repeat Unit 34 | A | D | S | S | I | D | Q | S | G | N | N | | N | L | A | D | V | T | Q | G | G | F | D | |
| Repeat Unit 35 | N | T | S | T | I | V | Q | S | G | N | N | | N | A | A | T | V | N | Q | S | S | T | G | |
| Repeat Unit 36 | N | V | S | D | I | I | Q | S | G | N | G | | N | S | A | T | V | T | Q | G | G | G | V | I |

| EI43-CsgA | β1 strand | | | | | | | Intra-Repeat Loops | | | | | β2 strand | | | | | | | Inter-Repeat Loops | | | | |
|---|---|---|---|---|---|---|---|---|---|---|---|---|---|---|---|---|---|---|---|---|---|---|---|---|
| consensus motif | N | Ω | Ψ | Ω | Ψ | Ω | Q | x | G | x | x | Σ | n | Ω | Ψ | Ω | Ψ | Ω | Q | x | G | x | x | Σ |
| N-terminal motif | M | R | L | W | A | R | Q | T | K | R | S | SAGVLGEILMKKTA | L | L | G | V | S | I | I | A | L | S | A | AAPAFGQ |
| Repeat Unit 1 | S | Q | S | T | V | T | Q | T | G | N | N | | S | T | V | D | V | T | Q | G | G | P | D | GG |
| Repeat Unit 2 | N | V | S | T | V | D | Q | S | A | S | D | | S | N | V | T | I | N | Q | E | G | F | D | DNLSGVT |
| Repeat Unit 3 | N | T | S | T | V | T | Q | A | G | L | N | | Q | D | V | Q | S | T | Q | V | G | D | D | |
| Repeat Unit 4 | Q | T | S | S | V | I | Q | S | G | V | D | | M | E | A | L | L | V | Q | G | G | A | G | |
| Repeat Unit 5 | N | S | S | T | I | N | Q | S | D | T | D | | N | F | A | D | V | I | Q | D | G | D | D | |
| Repeat Unit 6 | N | I | S | N | V | S | Q | S | D | E | D | | G | T | V | D | V | D | Q | L | G | E | R | L |
| Repeat Unit 7 | T | S | T | I | S | Q | Q | D | D | N | N | | Q | N | A | V | V | L | Q | T | N | A | D | |
| Repeat Unit 8 | N | T | S | T | I | T | Q | R | A | S | N | | S | D | V | F | V | T | Q | S | G | A | E | |
| Repeat Unit 9 | N | T | S | T | V | L | Q | G | N | I | S | DN | Q | E | A | T | I | L | Q | T | G | D | R | |
| Repeat Unit 10 | N | S | S | D | V | Q | Q | G | F | D | L | SGPLDDD | N | V | T | F | V | T | Q | N | G | D | D | |
| Repeat Unit 11 | N | T | A | V | V | R | Q | I | N | E | R | | N | L | A | D | I | L | Q | E | G | D | R | N |
| Repeat Unit 12 | E | A | R | V | V | N | Q | G | T | G | V | SGGGNED | N | I | A | D | I | D | Q | F | G | D | D | |
| Repeat Unit 13 | N | F | A | S | I | N | Q | P | G | E | D | | G | T | F | T | V | V | Q | N | G | F | D | |
| Repeat Unit 14 | S | S | S | T | G | L | Q | S | G | S | N | | D | T | A | S | V | I | Q | N | G | E | F | |
| Repeat Unit 15 | D | S | S | N | V | E | Q | S | G | S | G | | N | N | A | T | V | T | Q | E | G | F | D | LDGSLLA |
| Repeat Unit 16 | N | S | S | T | V | L | Q | S | G | T | G | | D | T | A | T | V | S | Q | V | G | D | N | |
| Repeat Unit 17 | Q | S | S | F | V | A | Q | S | G | S | V | | S | T | A | S | V | V | Q | G | G | V | G | |
| Repeat Unit 18 | N | S | S | G | I | T | Q | S | G | T | D | | V | T | A | T | V | N | Q | E | G | D | G | |
| Repeat Unit 19 | N | S | S | T | V | T | Q | A | N | S | L | | S | D | A | V | V | S | Q | T | G | D | F | |
| Repeat Unit 20 | D | T | S | I | V | A | Q | S | G | I | D | | E | Q | A | N | V | D | Q | A | G | E | T | |
| Repeat Unit 21 | G | V | S | E | V | T | Q | S | G | D | S | | N | F | A | D | V | F | Q | R | G | A | E | |
| Repeat Unit 22 | N | T | S | F | I | T | Q | S | G | L | D | | G | N | V | D | L | D | Q | T | G | D | E | |
| Repeat Unit 23 | N | F | S | N | A | I | Q | A | G | F | E | | G | D | I | I | V | N | Q | T | G | D | G | |
| Repeat Unit 24 | N | S | S | I | V | N | Q | Q | G | D | A | TGF | N | F | A | T | I | D | Q | S | G | N | G | |
| Repeat Unit 25 | N | T | S | E | A | I | Q | F | N | N | S | | N | N | A | N | I | L | Q | S | G | D | D | |
| Repeat Unit 26 | N | T | S | F | A | T | Q | T | G | T | G | | V | I | A | D | V | D | Q | I | G | E | G | |
| Repeat Unit 27 | A | T | S | T | V | T | Q | D | G | S | G | | N | L | A | I | V | S | Q | E | G | I | D | FDLSGTT |
| Repeat Unit 28 | D | T | S | L | V | A | Q | T | G | T | G | | Q | N | A | E | V | T | Q | V | G | D | G | |
| Repeat Unit 29 | N | S | S | S | V | V | Q | N | N | A | D | | N | T | A | S | V | L | T | A | G | I | G | |
| Repeat Unit 30 | N | S | S | F | V | T | Q | N | G | L | T | | G | L | A | T | V | D | Q | Q | G | D | G | |
| Repeat Unit 31 | N | S | S | I | V | S | Q | G | G | S | N | | A | S | A | D | V | S | Q | Q | S | D | T | |
| Repeat Unit 32 | S | S | S | N | V | N | Q | A | G | D | D | | V | L | A | L | V | D | Q | A | G | A | N | |
| Repeat Unit 33 | H | S | A | S | I | V | Q | L | T | T | G | PTGGVDLN | N | G | A | L | I | D | Q | Q | G | D | G | |
| Repeat Unit 34 | N | T | A | N | I | T | Q | S | G | G | I | VFFNVLSGTGLSSG | Q | I | A | E | I | S | Q | N | G | S | G | |
| Repeat Unit 35 | N | T | G | T | V | A | Q | S | G | T | T | | N | I | G | R | F | F | Q | D | G | D | D | |
| Repeat Unit 36 | N | S | A | T | I | S | Q | T | G | Q | D | | S | D | S | S | F | G | Q | T | G | N | N | |
| Repeat Unit 37 | N | T | A | N | V | T | Q | A | L | I | A | PPGGI | A | F | A | D | P | N | Q | N | G | D | G | |
| Repeat Unit 38 | N | S | A | T | V | V | Q | N | G | T | V | TGFFT | T | R | A | Q | S | A | Q | I | G | N | G | |
| Repeat Unit 39 | N | T | L | T | T | T | Q | S | G | T | D | | D | I | V | F | I | D | Q | I | G | D | G | |
| Repeat Unit 40 | N | M | S | M | V | N | Q | L | A | G | G | VS | N | E | G | D | I | N | Q | T | G | D | D | |
| Repeat Unit 41 | G | I | S | E | L | T | Q | S | G | T | D | | Q | F | A | E | L | F | Q | S | G | D | L | |
| Repeat Unit 42 | N | T | S | L | I | T | Q | A | G | V | T | | N | T | A | T | V | T | Q | G | S | D | G | |
| Repeat Unit 43 | N | F | S | S | V | N | Q | N | G | T | G | | N | S | T | T | V | T | Q | | | | | |

| Er46-CsgA | β1 strand | | | | | | | Intre-Repeat Loops | | | | | β2 strand | | | | | | | Inter-Repeat Loops | | | | |
|---|---|---|---|---|---|---|---|---|---|---|---|---|---|---|---|---|---|---|---|---|---|---|---|---|
| | n | Ω | Ψ | Ω | Ψ | Ω | Q | x | G | x | x | Σ | N | Ω | Ψ | Ω | Ψ | Ω | Q | x | G | x | x | Σ |
| N-terminal motif | M | K | F | S | A | R | K | T | L | L | L | TGVAVIAM | S | A | A | A | S | A | Q | E | A | | | |
| Repeat Unit 1 | N | T | G | Y | I | G | Q | R | D | D | S | | N | F | A | Y | L | S | Q | E | G | A | | |
| Repeat Unit 2 | N | R | G | F | I | Y | Q | H | G | Q | Q | | N | V | F | L | P | L | A | A | G | G | A | AGDAQQGK |
| Repeat Unit 3 | N | D | L | G | V | M | Q | L | G | S | K | NV | V | G | G | D | F | L | Q | K | G | E | | |
| Repeat Unit 4 | N | V | G | G | I | A | Q | F | G | T | N | N | L | V | D | K | F | R | Q | D | G | A | | |
| Repeat Unit 5 | N | R | G | T | I | F | Q | G | G | N | A | | N | V | A | R | N | T | Q | E | E | G | D | SG |
| Repeat Unit 6 | N | D | T | G | I | I | Q | L | N | D | S | PNKPG | N | Y | A | E | S | N | Q | L | G | A | N | GG |
| Repeat Unit 7 | N | F | S | G | I | L | Q | N | G | S | G | | N | A | A | I | N | N | Q | T | D | E | T | S |
| Repeat Unit 8 | S | N | A | V | T | A | Q | S | G | T | G | | N | A | A | I | N | N | Q | T | A | S | I | D |
| Repeat Unit 9 | G | N | A | G | I | L | Q | V | G | K | G | | N | G | A | I | N | T | Q | T | G | S | S | LD |
| Repeat Unit 10 | A | T | A | I | I | A | Q | N | G | K | D | | N | S | A | V | N | T | Q | T | N | V | D | P |
| Repeat Unit 11 | T | S | A | T | V | I | Q | D | G | N | N | | N | A | A | A | N | V | Q | T | S | V | N | N |
| Repeat Unit 12 | S | N | A | F | I | L | Q | D | G | N | N | | N | T | A | S | N | L | Q | S | P | G | T | QAGPADS |
| Repeat Unit 13 | L | N | A | S | I | A | Q | S | G | N | K | | N | N | A | V | N | L | Q | A | G | G | S | GG |
| Repeat Unit 14 | G | K | K | N | T | A | Q | T | V | Q | D | GNR | N | T | A | L | N | S | Q | I | G | D | Y | HNNGYNND |
| Repeat Unit 15 | N | K | A | V | I | G | Q | V | G | N | D | | N | L | A | V | N | T | Q | N | D | G | R | N |
| Repeat Unit 16 | S | T | A | T | I | L | Q | G | G | D | R | | N | D | A | Y | N | L | S | E | G | G | T | N |
| Repeat Unit 17 | H | S | S | A | I | A | Q | F | G | D | D | | N | L | A | V | N | A | R | N | R | G | G | NPIEG |
| Repeat Unit 18 | N | D | A | A | I | L | Q | F | G | D | D | | N | I | A | A | N | Y | Q | N | D | A | I | GSEQFTN |
| Repeat Unit 19 | N | S | A | N | I | G | Q | F | G | S | G | | N | I | A | T | N | F | Q | D | T | G | S | D |
| Repeat Unit 20 | N | E | G | N | I | T | Q | F | G | T | N | | N | G | A | S | N | A | Q | V | G | G | G | VE |
| Repeat Unit 21 | N | T | A | N | I | L | Q | I | G | S | G | | N | S | G | A | N | V | Q | S | G | G | T | E |
| Repeat Unit 22 | A | T | A | N | I | V | Q | G | G | S | N | | N | V | A | A | N | L | Q | T | N | V | D | P |
| Repeat Unit 23 | T | N | A | S | I | L | Q | L | G | S | R | | N | T | A | G | N | V | Q | A | D | L | V | N |
| Repeat Unit 24 | S | D | A | L | I | V | Q | D | G | A | R | | N | E | A | S | N | L | Q | T | D | G | T | N |
| Repeat Unit 25 | M | A | T | V | I | V | Q | K | G | D | R | | N | N | A | G | T | I | Q | A | G | S | T | D |
| Repeat Unit 26 | V | D | A | F | T | L | Q | N | G | D | R | | N | E | A | S | N | G | Q | S | G | V | T | G |
| Repeat Unit 27 | G | T | A | G | I | I | Q | D | G | N | R | | N | V | A | G | S | V | Q | L | T | S | S | D |
| Repeat Unit 28 | V | D | A | S | I | I | Q | S | G | N | R | | N | D | A | A | N | L | Q | S | G | V | T | G |
| Repeat Unit 29 | G | S | A | G | I | L | Q | D | G | R | D | | N | T | A | A | N | L | Q | L | S | S | T | G |
| Repeat Unit 30 | V | D | A | T | I | I | Q | S | G | R | F | | N | E | A | V | N | S | Q | T | A | S | S | L |
| Repeat Unit 31 | S | N | A | F | I | I | Q | G | A | G | G | N | N | E | S | A | N | V | Q | D | D | V | S | G |
| Repeat Unit 32 | A | T | A | L | I | G | Q | F | G | W | G | | N | T | A | G | N | L | Q | Q | A | D | S | SN |
| Repeat Unit 33 | V | I | A | G | I | L | Q | V | G | S | G | | N | T | A | Y | N | T | Q | G | A | T | Q | TATYEVLPAF |
| Repeat Unit 34 | N | L | T | T | T | A | S | V | P | G | L | SN | Y | L | T | P | N | M | Q | G | P | F | I | FGIAGGPTTLTGSGSAPAED |
| Repeat Unit 35 | S | I | A | L | V | A | Q | F | G | N | D | | N | S | A | A | N | V | Q | I | G | Q | L | IAPASGIASITTTTY |
| Repeat Unit 36 | g | i | n | t | s | v | Q | n | g | g | a | SWIIGPKT | i | t | h | s | q | a | Q | s | t | q | t | t |
| Repeat Unit 37 | S | t | p | f | t | n | p | l | g | e | i | | N | v | p | t | v | a | | | | | | |
| Repeat Unit 38 | S | A | A | G | I | V | Q | V | G | D | D | | N | D | A | F | N | F | Q | T | V | A | A | G |
| Repeat Unit 39 | S | L | A | A | T | V | Q | I | G | D | G | | N | G | A | A | N | V | Q | T | A | T | L | G |
| Repeat Unit 40 | A | A | A | A | I | V | Q | S | G | N | G | | N | T | A | S | N | L | Q | N | G | A | I | L |
| Repeat Unit 41 | S | A | A | G | I | I | Q | A | G | H | D | | N | V | A | T | T | T | Q | I | A | S | L | S |
| Repeat Unit 42 | N | A | A | L | T | I | Q | N | G | K | K | | N | V | A | G | T | N | Q | S | N | T | V | G |
| Repeat Unit 43 | N | T | S | L | I | V | Q | S | S | N | N | | S | L | A | L | V | D | Q | T | G | A | G | G |
| Repeat Unit 44 | H | S | S | A | V | V | Q | A | G | A | Y | | S | N | A | W | I | S | Q | T | G | S | A | |
| Repeat Unit 45 | S | N | S | V | V | L | Q | Y | G | S | G | SSDADR | N | Y | A | I | V | S | Q | G | N | S | G | |
| Repeat Unit 46 | Q | N | S | V | V | I | Q | A | G | R | A | | N | V | A | F | V | S | Q | N | | | | |

**Supplementary Figure 4. Top view of β-solenoid proteins.** Interloop distances of the β-solenoid proteins.

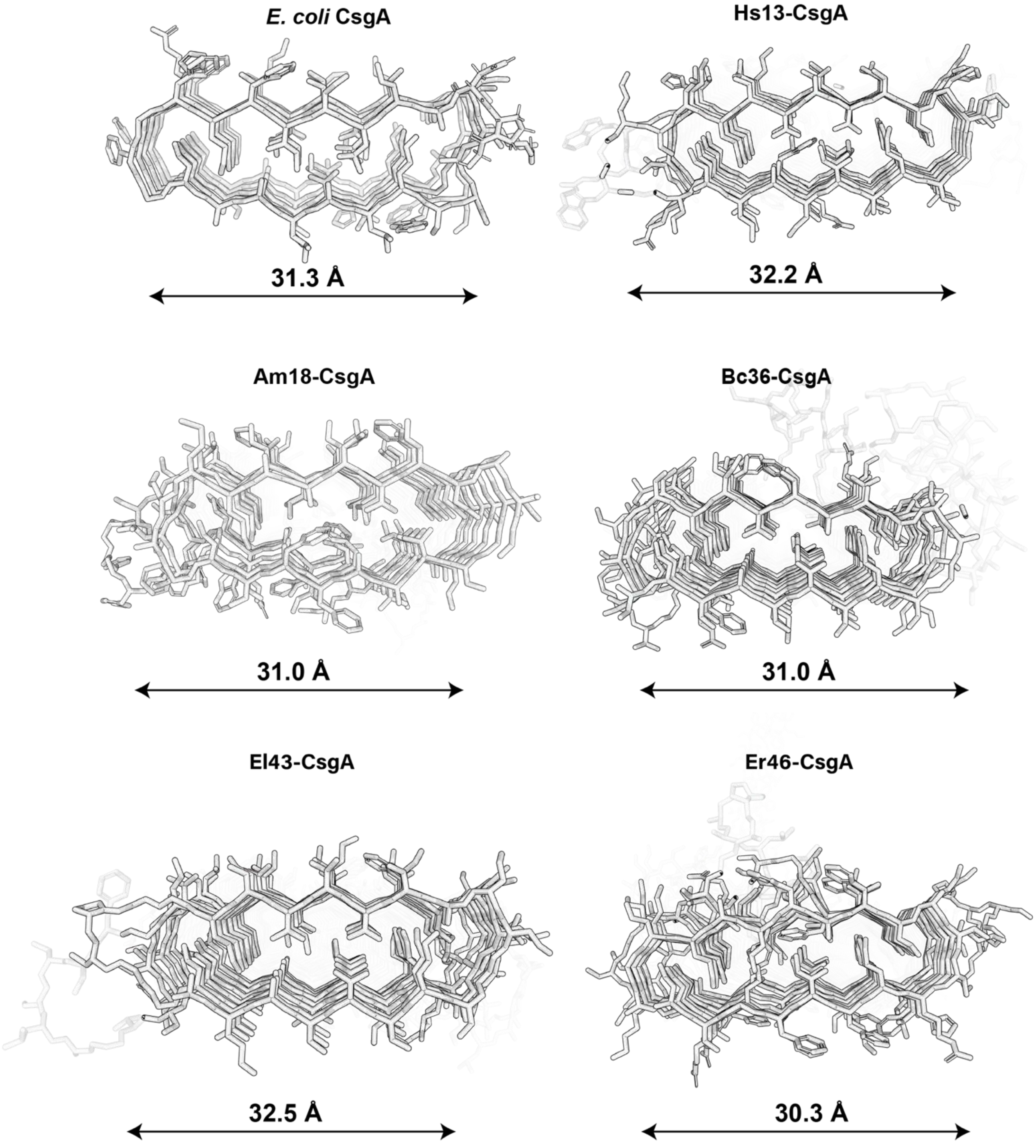

**Supplementary Table 1. Amino acid composition of the β-solenoid proteins.** The list shows the percentage of amino acid residues in CsgA, Hs13-CsgA, Am18-CsgA, Bc36-CsgA, El43-CsgA and Er46-CsgA.

| Amino acid Residue % | CsgA | Hs13-CsgA | Am18-CsgA | Bc36-CsgA | El43-CsgA | Er46-CsgA |
|---|---|---|---|---|---|---|
| ***G*** | 21.37 | 12.22 | 10.9 | 13.22 | 12.96 | 13.54 |
| ***N*** | 12.21 | 10.14 | 11.21 | 9.97 | 9.95 | 13.54 |
| ***S*** | 9.16 | 7.04 | 11.53 | 11.81 | 12.11 | 9.03 |
| ***Q*** | 8.4 | 4.97 | 9.97 | 8.99 | 9.95 | 8.52 |
| ***A*** | 7.63 | 8.28 | 8.1 | 10.73 | 7.23 | 13.2 |
| ***T*** | 6.87 | 6.42 | 6.85 | 13.76 | 11.08 | 8.09 |
| ***V*** | 6.11 | 10.77 | 7.79 | 8.88 | 8.54 | 6.13 |
| ***D*** | 6.11 | 8.7 | 7.17 | 7.37 | 9.58 | 5.28 |
| ***L*** | 4.58 | 6.42 | 4.98 | 2.6 | 4.51 | 5.45 |
| ***Y*** | 3.05 | 1.24 | 3.12 | 0.87 | 0 | 1.28 |
| ***I*** | 2.29 | 5.8 | 3.43 | 4.01 | 4.88 | 5.79 |
| ***F*** | 2.29 | 4.14 | 2.49 | 1.95 | 3.47 | 2.39 |
| ***H*** | 2.29 | 0.21 | 2.18 | 0 | 0.09 | 0.51 |
| ***P*** | 1.53 | 0.62 | 0.31 | 1.95 | 0.75 | 1.62 |
| ***K*** | 1.53 | 1.04 | 2.49 | 0.43 | 0.28 | 1.45 |
| ***R*** | 1.53 | 0.62 | 1.56 | 1.3 | 1.22 | 1.79 |
| ***E*** | 1.53 | 7.87 | 2.8 | 1.73 | 2.82 | 1.7 |
| ***M*** | 0.76 | 2.28 | 2.18 | 0.43 | 0.47 | 0.43 |
| ***W*** | 0.76 | 1.24 | 0.93 | 0 | 0.09 | 0.26 |
| ***C*** | 0 | 0 | 0 | 0 | 0 | 0 |

**Supplementary Table 2. Physiochemical properties of β-solenoid proteins.** The list shows the calculated/estimated molecular weight, isoelectric point, net charge, surface area (SASA) and hydrophobicity (Gravy) of CsgA, Hs13-CsgA, Am18-CsgA, Bc36-CsgA, El43-CsgA and Er46-CsgA.

| PROTEIN | Molecular Weight (kDa) | Isoelectric Point (Pi) | Net Charge at Ph 7.4 (Z) | Interpretation | SASA (A^2) | Gravy |
|---|---|---|---|---|---|---|
| **CsgA** | 13.1 | 4.528 | -6.319 | Slightly acidic; minor negative charge at pH 7.4, moderately stable and soluble. | 13247.397 | -0.7183 |
| **Hs13-CsgA** | 34.02 | 4.401 | -19.261 | Highly acidic; significantly negative charge at pH 7.4, highly soluble, may strongly repel other negatively charged molecules. | 20327.082 | -0.6374 |
| **Am18-CsgA** | 50.59 | 3.358 | -72.658 | Slightly acidic; moderate negative charge at pH 7.4, stable and soluble. | 15050.855 | -0.1058 |
| **Bc36-CsgA** | 91.04 | 3.519 | -68.719 | Highly acidic; significantly negative charge at pH 7.4, highly soluble, may strongly repel other negatively charged molecules. | 37039.582 | -0.4354 |
| **El43-CsgA** | 107.81 | 3.325 | -116.609 | Extremely acidic; very high negative charge at pH 7.4, extremely soluble, strong repulsion likely. | 105950.086 | -0.4524 |
| **Er46-CsgA** | 117.63 | 4.229 | -44.521 | Acidic; high negative charge at pH 7.4, high solubility and potential for repulsive interactions. | 117655.484 | -0.3599 |

**Supplementary Figure 5. Gravy hydrophobicity index of β-solenoid proteins.** The hydrophobicity of CsgA, Hs13-CsgA, Am18-CsgA, Bc36-CsgA, El43-CsgA and Er46-CsgA is presented along with their Gravy index.

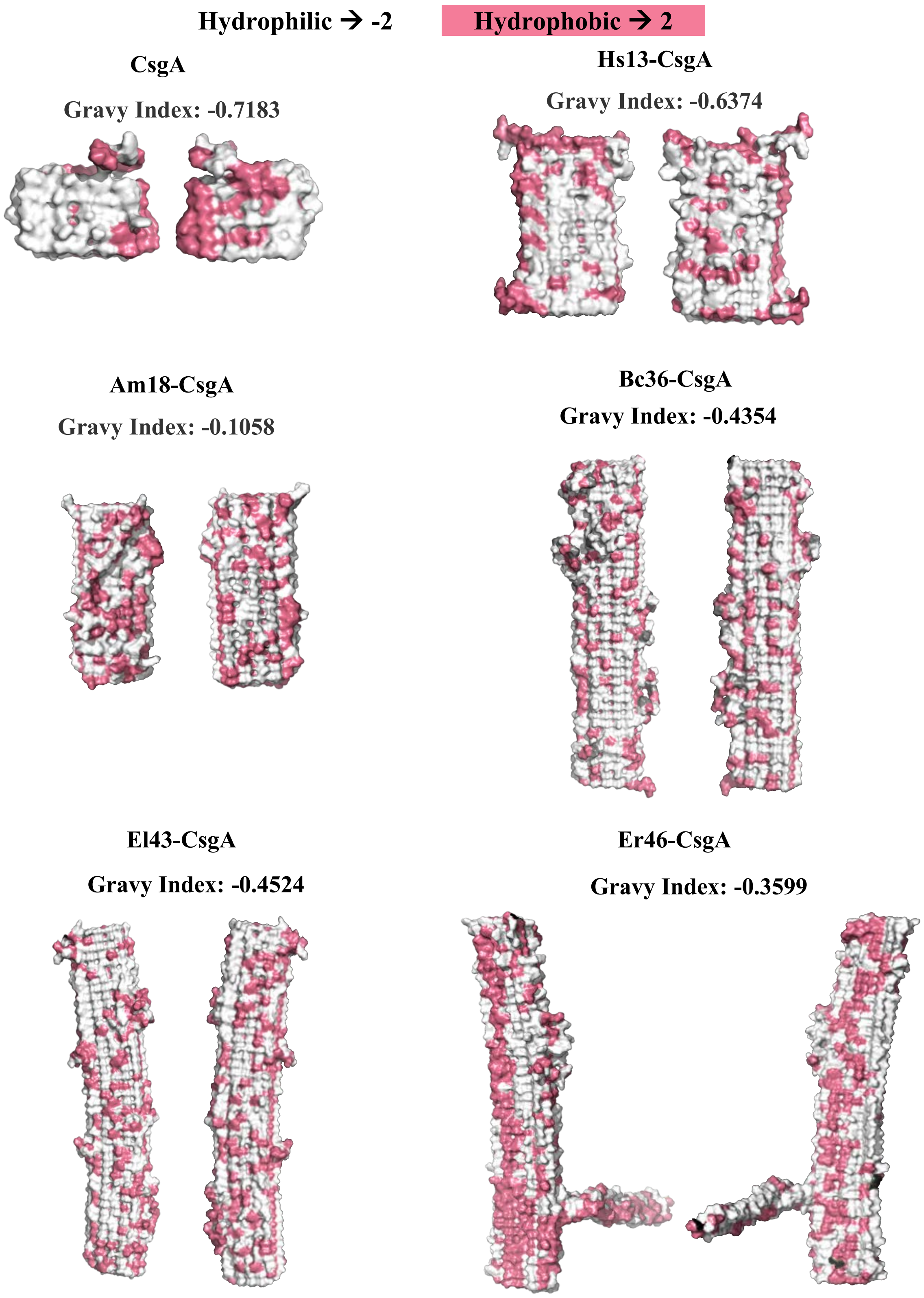

**Supplementary Figure 6. The structure of β-solenoid proteins predicted by AlphaFold2.** Structure of CsgA, Hs13-CsgA, Am18-CsgA, Bc36-CsgA, El43-CsgA and Er46-CsgA predicted by AlphaFold2 along with their pLDDT (predicted local distance difference test) scores.

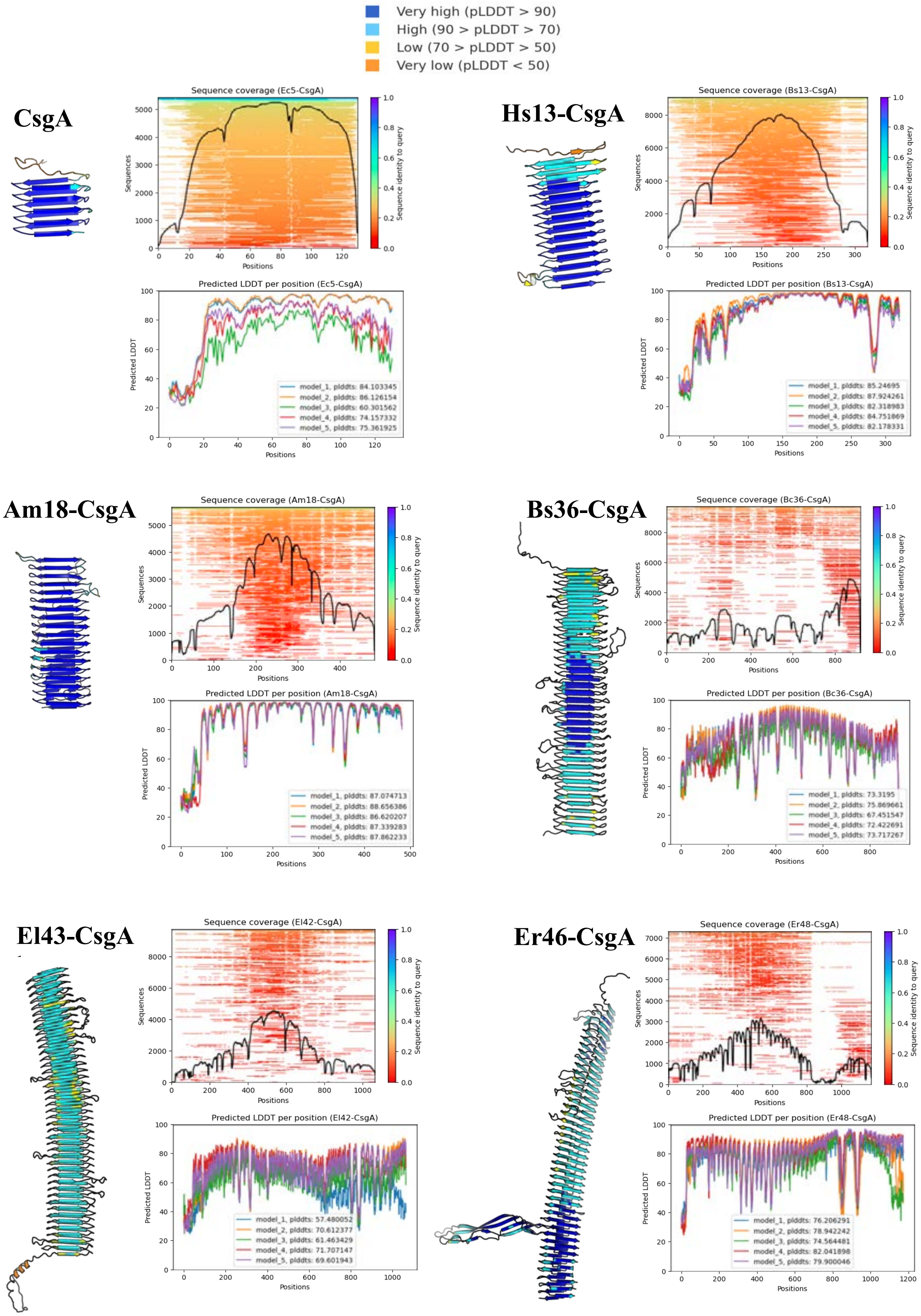

**Supplementary Figure 7. Comparison of β-solenoid protein structures obtained using AlphaFold2 and molecular dynamics simulation.** The structures of CsgA, Hs13-CsgA, Am18-CsgA, Bc36-CsgA, El43-CsgA, and Er46-CsgA predicted using AlphaFold2 and molecular dynamics simulation are overlaid to show their similarity. The corresponding pLDDT (predicted local distance difference test) scores for the entire β-solenoid protein and RMSD (root mean square deviation) values of the β-solenoid core (without outlier regions and N-terminal) are presented.

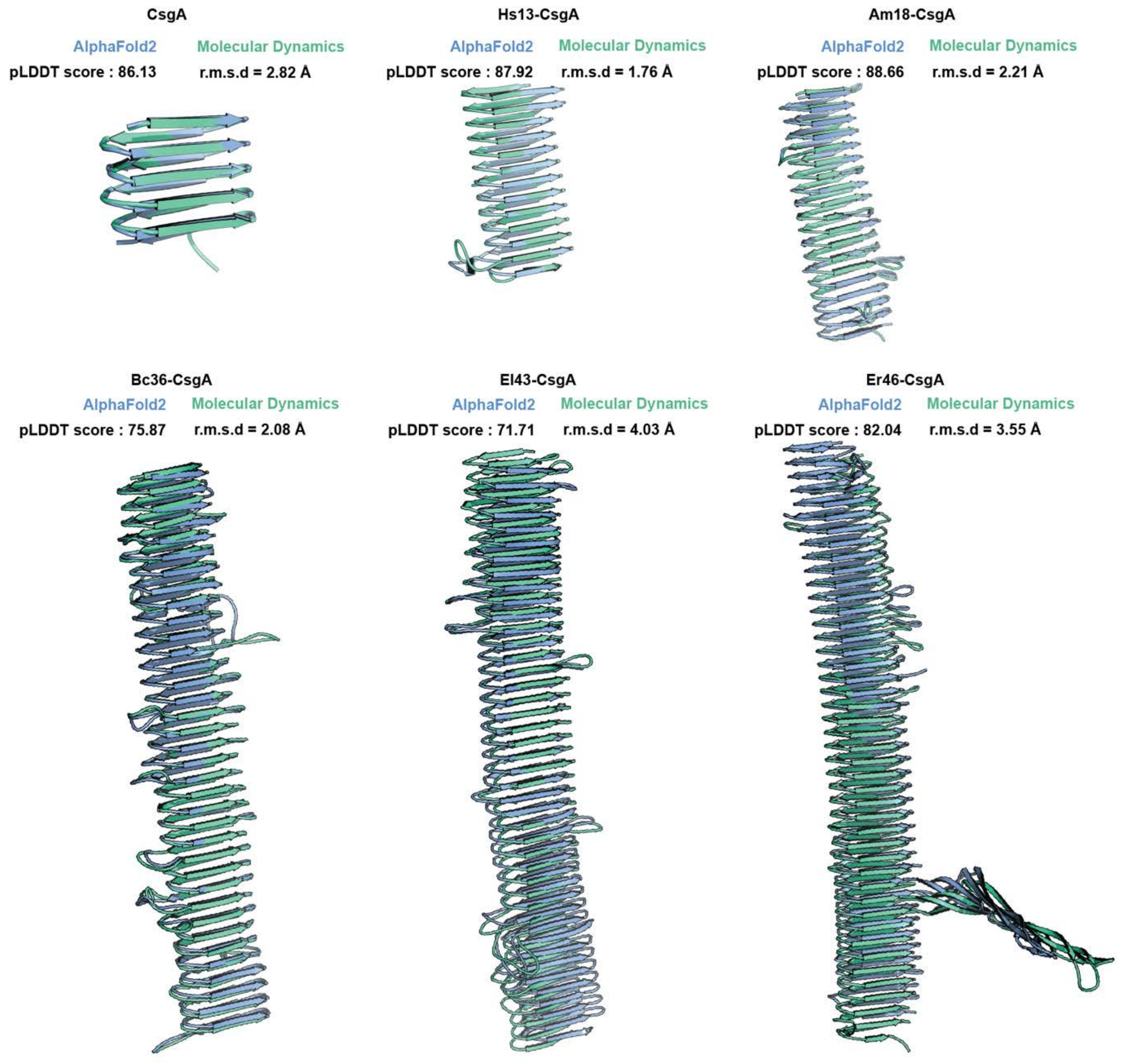

**Supplementary Figure 8: RMSD of the β-solenoid proteins**. Molecular dynamics simulation based RMSD (root mean square deviation) values for 150 ns of entire CsgA, Hs13-CsgA, Am18-CsgA, Bc36-CsgA, El43-CsgA and Er46-CsgA.

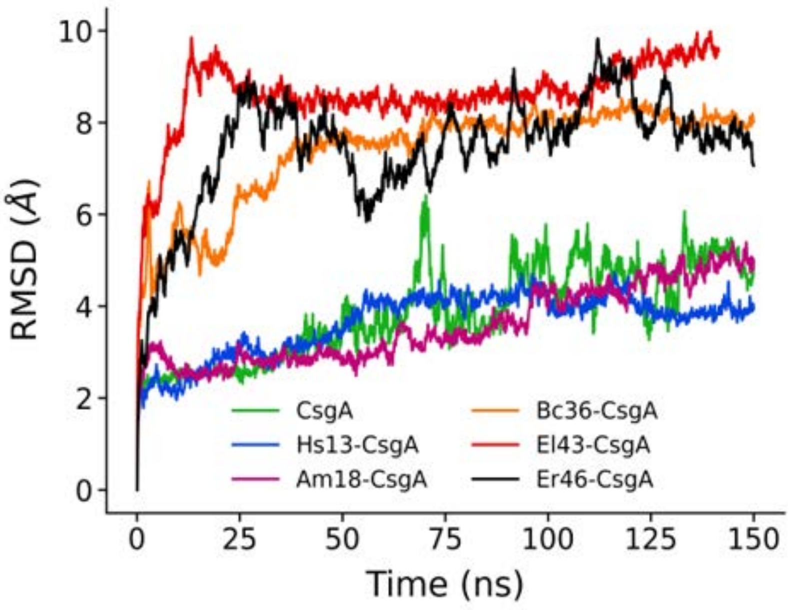

**Supplementary Figure 9: RMSF of the β-solenoid proteins.** Molecular dynamics simulation based RMSF (root mean squared fluctuations) values of entire CsgA, Hs13-CsgA, Am18-CsgA, Bc36-CsgA, El43-CsgA and Er46-CsgA. The dark lines show the average RMSF over the last 50 ns production run and the standard deviations are shown in lighter shades.

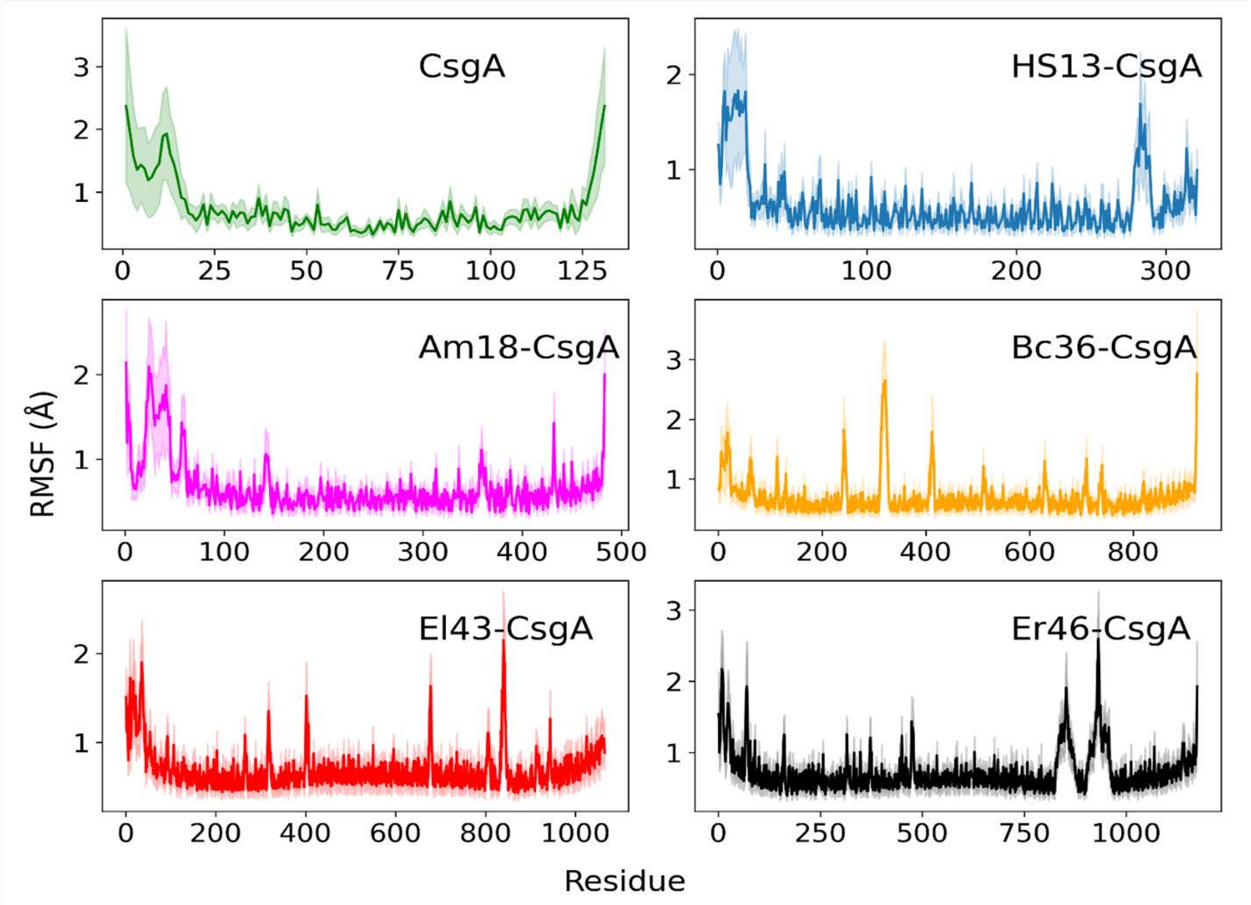

**Supplementary Figure 10. Ramachandran plots of β-solenoid proteins.** The Ramachandran plots show the dihedral angles $\phi$ and $\psi$ for CsgA, Hs13-CsgA, Am18-CsgA, Bc36-CsgA, El43-CsgA and Er46-CsgA, as a function of simulation time intervals.

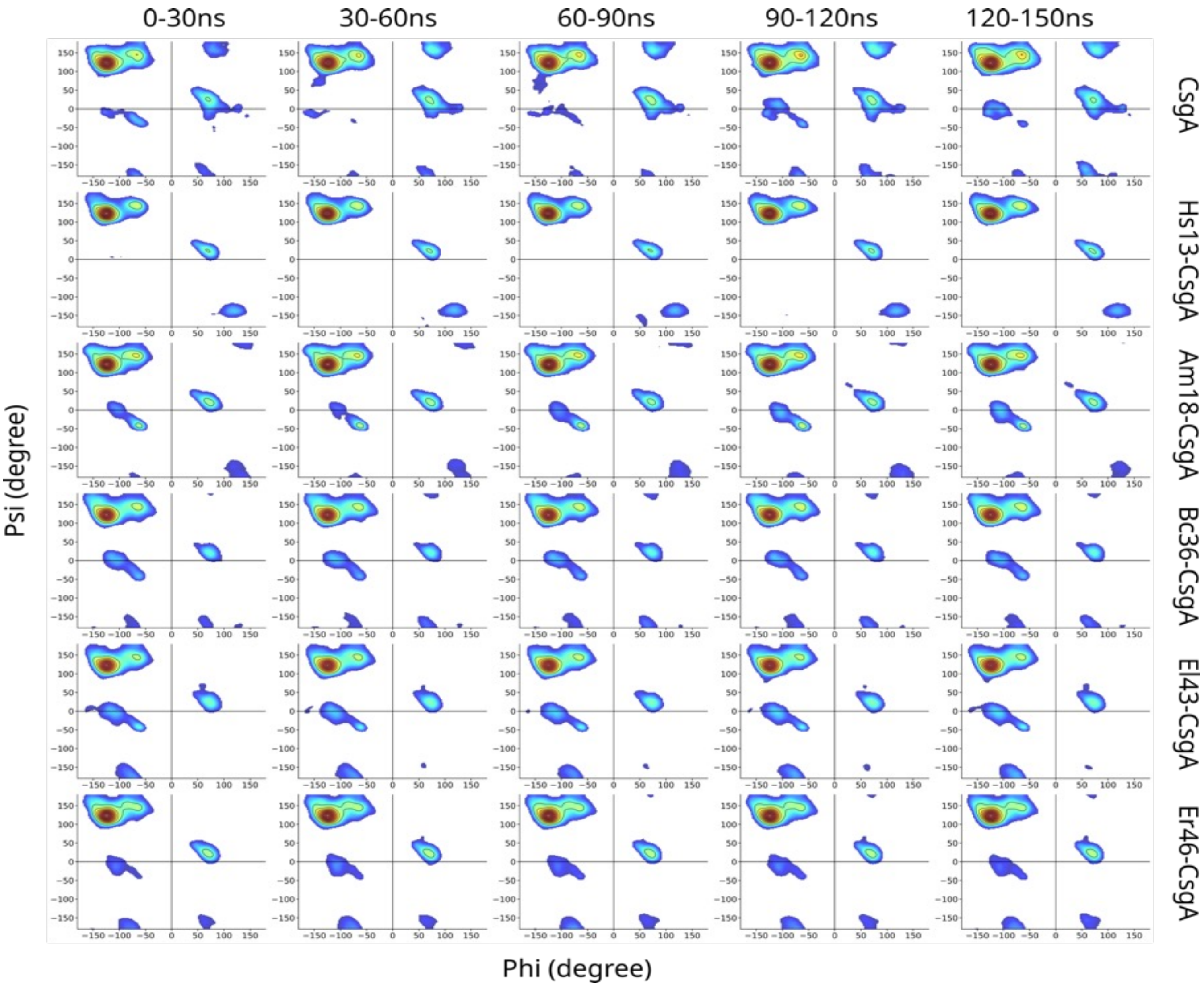

**Supplementary Table 3. Percentage of α-sheet and β-turns in β-solenoid proteins.**

| α-sheet & β-turn percentage | | |
|---|---|---|
| **Variants** | **α-sheet** | **β-turn** |
| **CsgA** | 3.37 | 18.69 |
| **Hs13-CsgA** | 2.90 | 13.58 |
| **Am18-CsgA** | 5.75 | 21.39 |
| **Bc36-CsgA** | 3.08 | 19.97 |
| **El43-CsgA** | 3.99 | 18.69 |
| **Er46-CsgA** | 3.13 | 15.88 |

**Supplementary Figure 11. Amino Acid composition of β-solenoid proteins.** The plots show the percentage of each amino acid present in CsgA, Hs13-CsgA, Am18-CsgA, Bc36-CsgA, El43-CsgA and Er46-CsgA. The positively charged residues (R and K) and the negatively charged residues (D and E) are highlighted in blue and red, respectively. The inset of each plot lists the total number of amino acid residues (N), net charge (Q), and the charge per residue (Q/N).

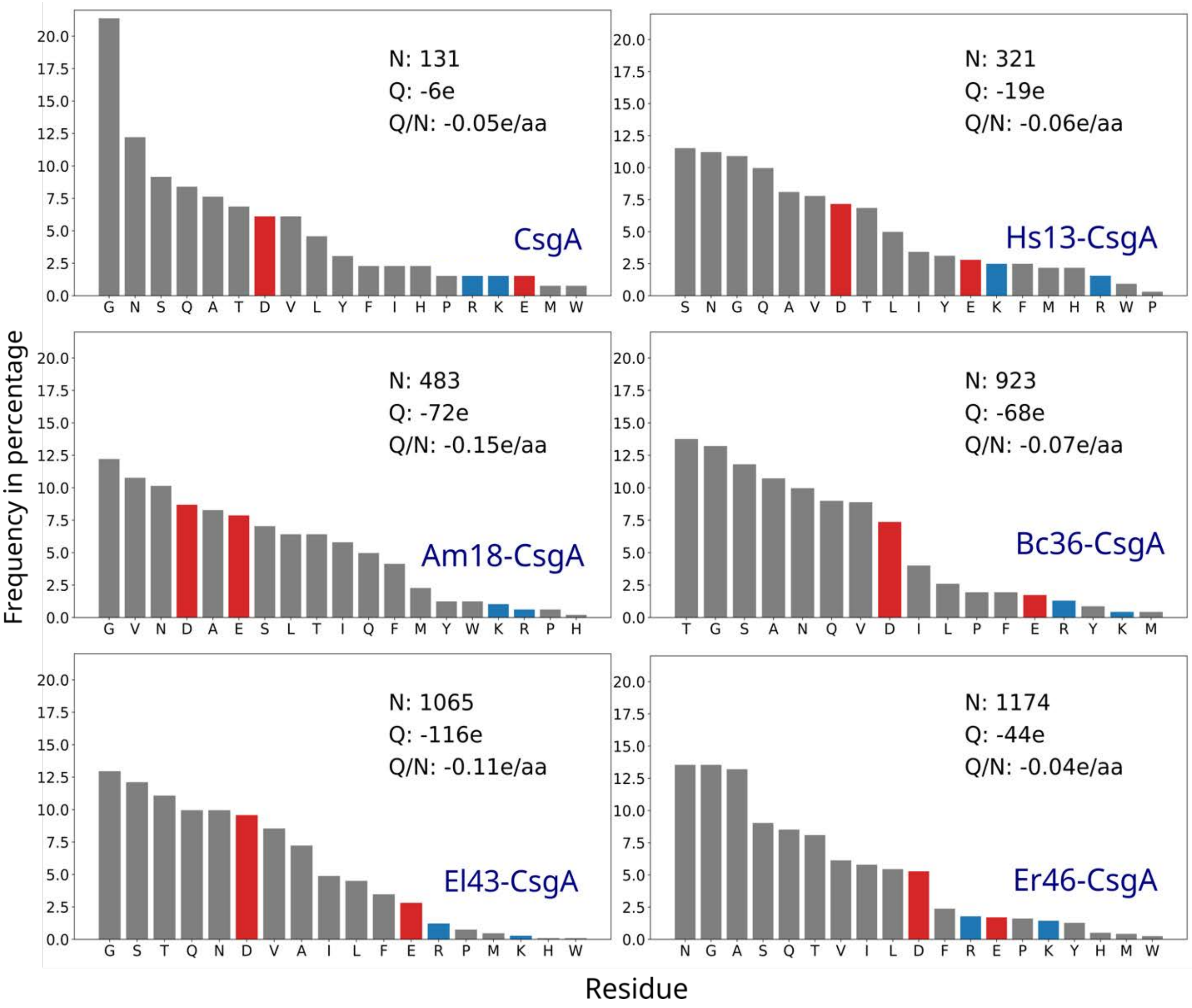

**Supplementary Figure 12. Heatmap of the average number of residues within the 12 Å surrounding of an amino acid residue.** The x-axis represents the amino acid residue in consideration, and the average surrounding residue counts are shown on the y-axis. The polar, non-polar, and charged residues are colored magenta, blue, and orange, respectively.

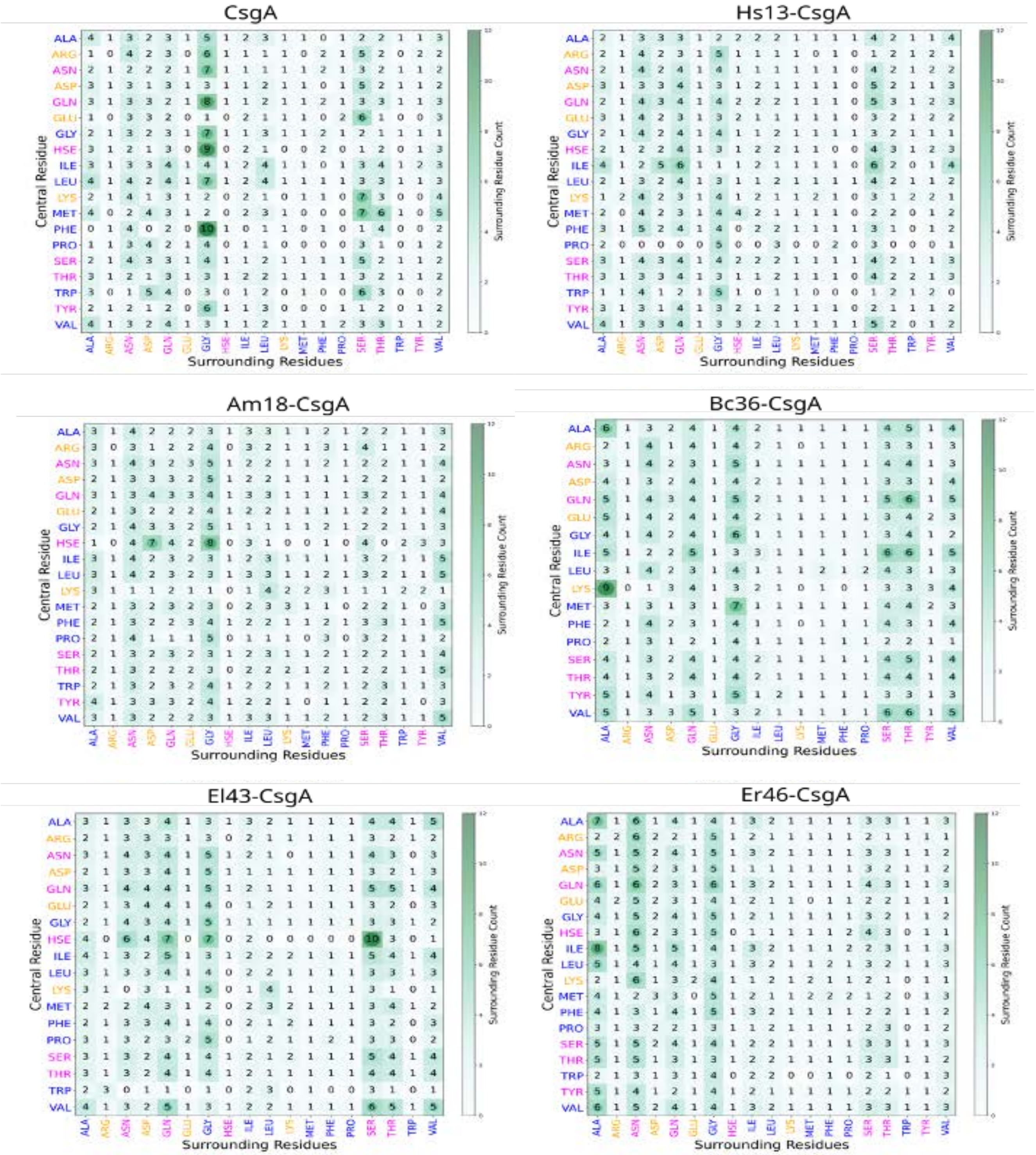

**Supplementary Figure 13. 3D printing of β-solenoid protein hydrogels.** The hydrogels of **a** Hs13-CsgA and **b** Er46-CsgA were utilized to show their utility for 3D printing. Scale bar 1 cm.

a

b

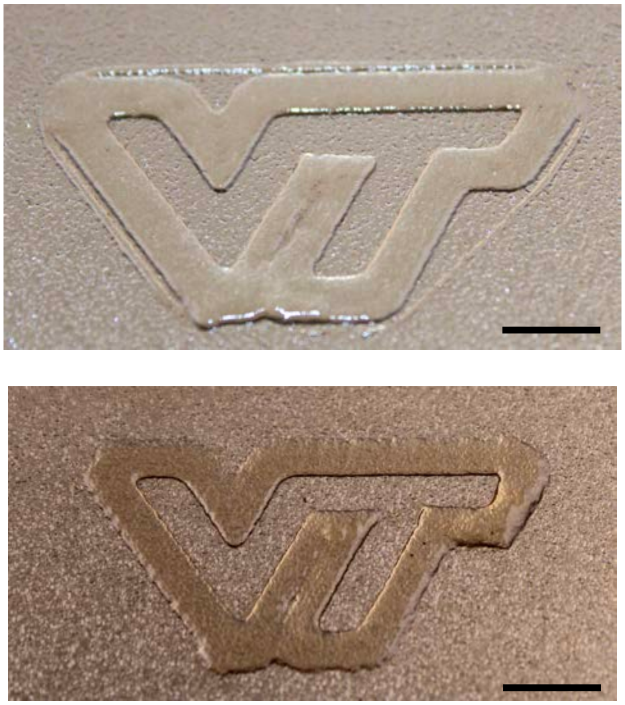

**Supplementary Figure 14. AlphaFold2 structure prediction of engineered β-solenoid proteins.** The structures of β-solenoid protein, Hs13-CsgA, that is genetically grafted with iron (Hs13-CsgA-IronBP) and antibody (Hs13-CsgA-IgG-BD) binding domains, predicted using AlphaFold2, is shown along with their pLDDT (predicted local distance difference test) scores.

**Hs13-CsgA-IronBP**

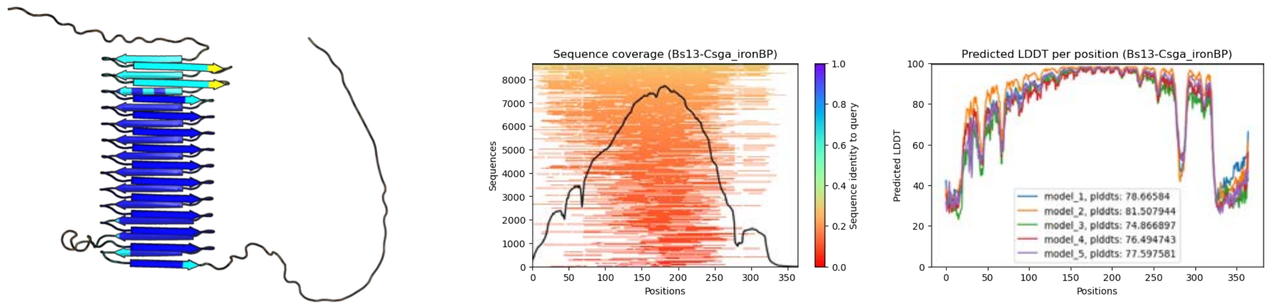


**Hs13-CsgA-IgG-BD**

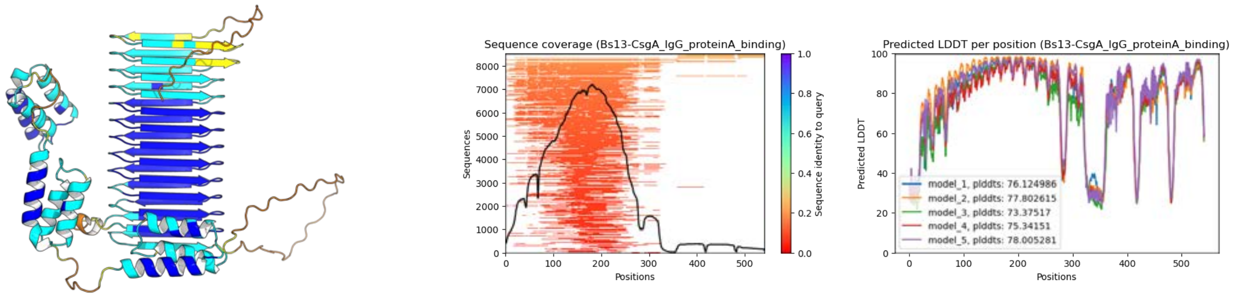

**Supplementary Figure 15. Congo Red Assay.** The plot shows the binding of Congo red dye to the engineered β-solenoid variants of Hs13-CsgA-IronBP and Hs13-CsgA-IgG-BD, indicating their cross-β characteristics. Biological replicates n = 4. Data represented as mean ± standard deviation. ****$p \leq 0.0001$, one-way ANOVA followed by Dunnett's test.

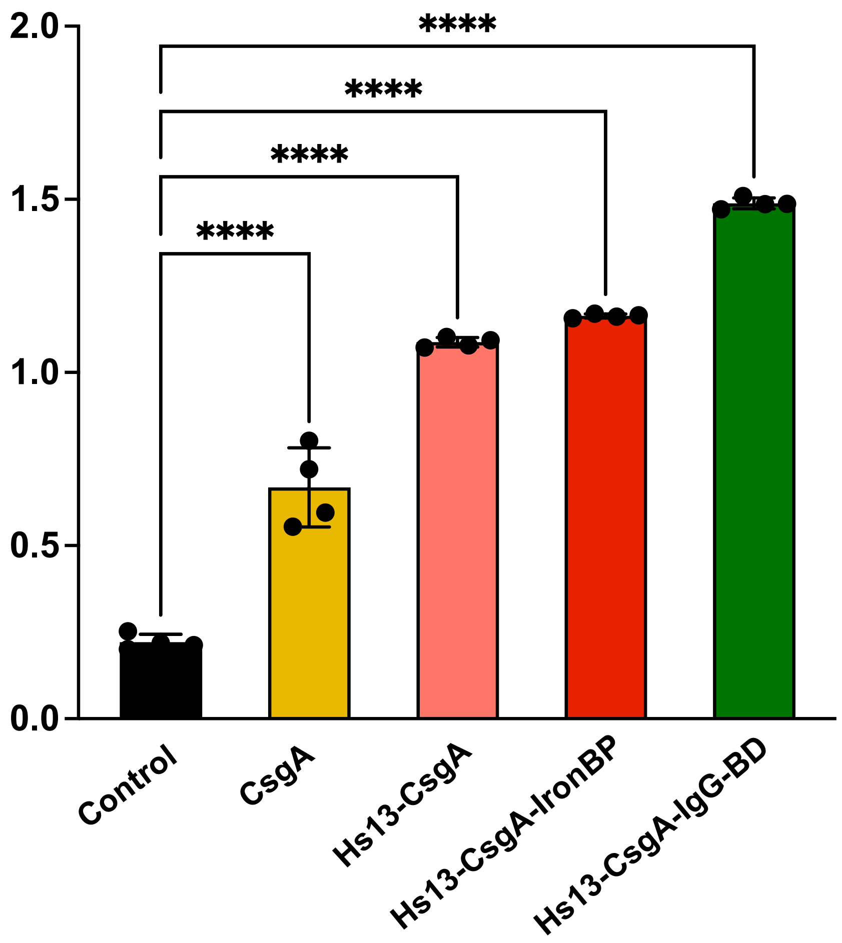

**Supplementary Figure 16. Energy dispersive X-ray analysis (EDAX) for elemental composition.** Representative EDAX plots of **a.** Hs13-CsgA and **b.** Hs13-CsgA-IronBP incubated with iron oxide nanoparticles show the elemental composition. Biological replicates n = 4.

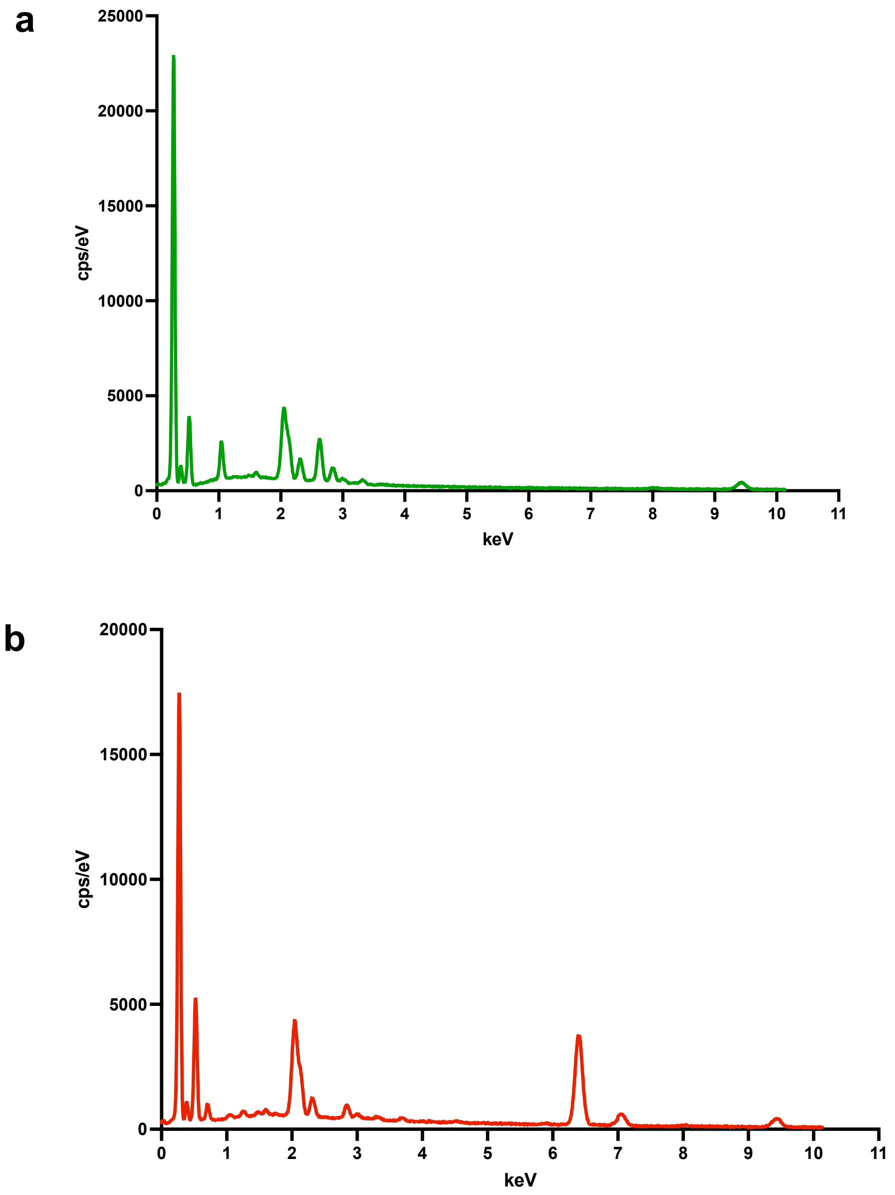

**Supplementary Figure 17. Plasmid map of β-solenoid protein variants used in this study.** The plasmid pET21d was separately cloned with genes of CsgA, Hs13-CsgA, Am18-CsgA, Bc36-CsgA, El43-CsgA, Er46-CsgA, Hs13-CsgA-IronBP and Hs13-CsgA-IgG-BD.

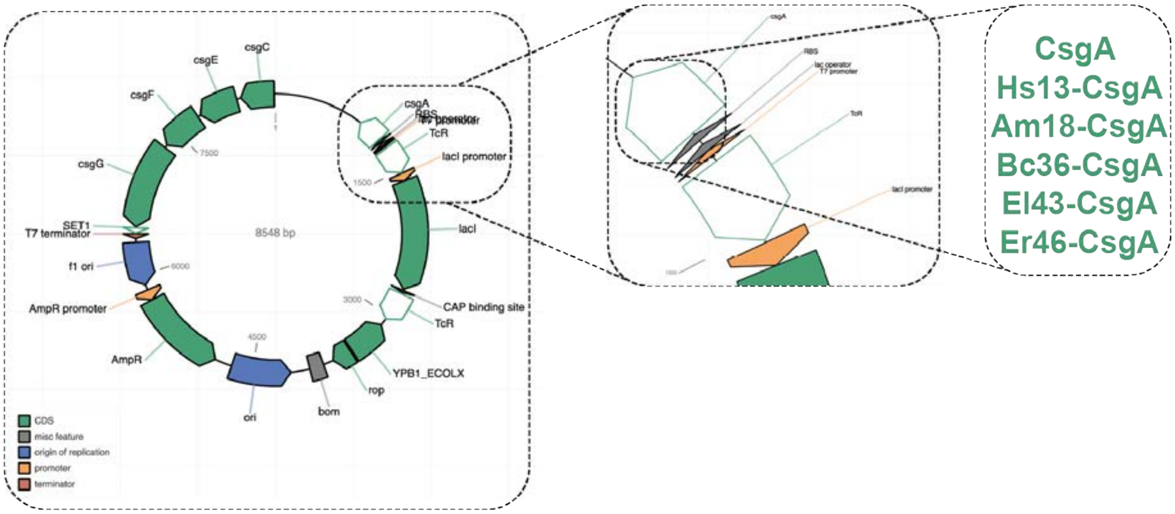


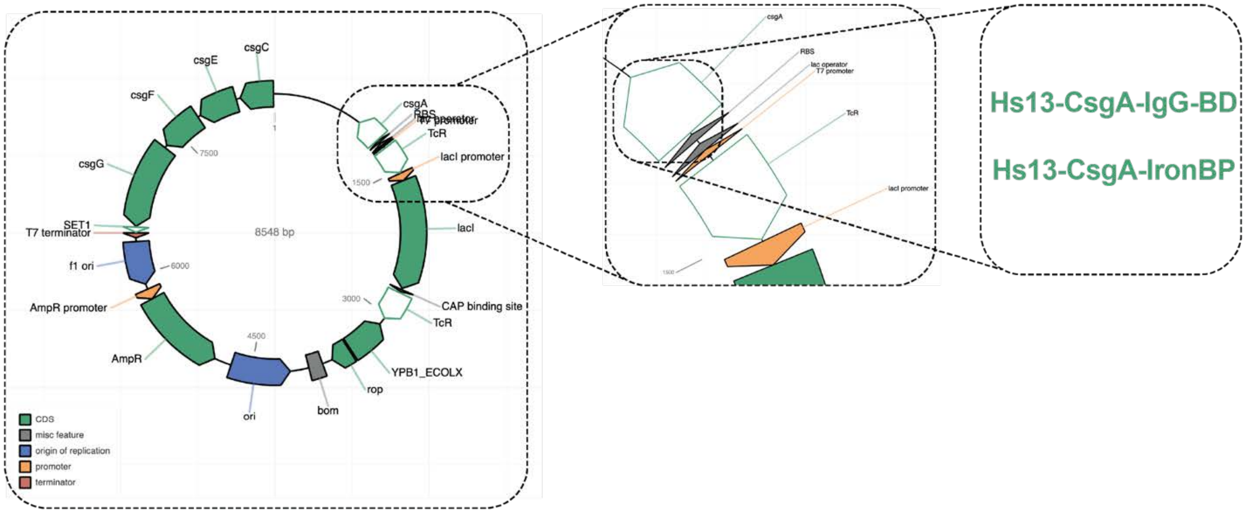

**Supplementary Table 4. Sequences of CsgA homologs and variants used in this study.** The list shows the protein and DNA sequences of CsgA, Hs13-CsgA, Am18-CsgA, Bc36-CsgA, El43-CsgA, Er46-CsgA, Hs13-CsgA-IronBP and Hs13-CsgA-IgG-BD.

| CsgA Homologs and Variants | Protein Sequence | DNA Sequence |
|---|---|---|
| **CsgA** | GVVPQYGGGGNHGGGGNNSGPNSELNIYQYG<br>GGNSALALQTDARNSDLTITQHGGGNGADVG<br>QGSDDSSIDLTQRGFGNSATLDQWNGKNSEM<br>TVKQFGGGNGAAVDQTASNSSVNVTQVGFGN<br>NATAHQY* | GGTGTTGTTCCTCAGTACGGCGGCGGCGGTAAC<br>CACGGTGGTGGCGGTAATAATAGCGGCCCAAAT<br>TCTGAGCTGAACATTTACCAGTACGGTGGCGGT<br>AACTCTGCACTTGCTCTGCAAACTGATGCCCGT<br>AACTCTGACTTGACTATTACCCAGCATGGCGGC<br>GGTAATGGTGCAGATGTTGGTCAGGGCTCAGAT<br>GACAGCTCAATCGATCTGACCCAACGTGGCTTC<br>GGTAACAGCGCTACTCTTGATCAGTGGAACGGC<br>AAAAATTCTGAAATGACGGTTAAACAGTTCGGT<br>GGTGGCAACGGTGCTGCAGTTGACCAGACTGCA<br>TCTAACTCCTCCGTCAACGTGACTCAGGTTGGC<br>TTTGGTAACAACGCGACCGCTCATCAGTACTAA |
| **Hs13-CsgA** | GVVPQYGGGGNHGGGGNNSGPNMKLKMTV<br>MAAAIAVSSTALANENESNIAQTGFGNDVSVT<br>QLTQGWGNNNSDVTQVGVNNGATVLQQGW<br>RGRNVSDIDQEGMGNNADALQHGTKNSFDLT<br>QSGMFNNSDSQQLGTGNALRVEQEGVKNNAT<br>TYQSGWRNYADISQSGSFNNSDVDQEGYLNV<br>AQVNQDGTKLSSDITQSGDYNFADVDQSGSFN<br>DSSIDQNGTQNKAEVLQSGLKNTSDIHQDGKL<br>NIAMVDQSGVNNDASIVQDGRYHQSEVTQSG<br>YNDVAYSSQSGYEHYSSITQSALPGFFGGLASL<br>ASNSATHVQMGVNNAAITTQVGGGNTAHVY<br>QH* | GGTGTTGTTCCTCAGTACGGCGGCGGCGGTAAC<br>CACGGTGGTGGCGGTAATAATAGCGGCCCAAAT<br>ATGAAGTTGAAGATGACGGTTATGGCCGCAGCT<br>ATCGCAGTTTCATCTACTGCCCTTGCAAATGAG<br>AATGAGTCAAATATCGCCCAAACTGGATTCGGT<br>AATGACGTCTCAGTTACACAATTAACGCAAGGT<br>TGGGGCAATAATAATTCAGACGTTACGCAAGTT<br>GGTGTCAATAATGGTGCAACAGTACTCCAACAA<br>GGTTGGAGAGGTCGGAATGTTAGTGACATAGAC<br>CAAGAGGGTATGGGAAATAATGCCGACGCTTTA<br>CAACACGGAACTAAGAATTCATTCGACCTCACT<br>CAAAGTGGAATGTTCAATAATTCTGACAGTCAA<br>CAACTTGGTACAGGTAATGCTTTACGTGTTGAG<br>CAAGAGGGAGTCAAGAATAATGCAACGACATA<br>CCAAAGTGGATGGCGTAATTACGCAGACATAA<br>GTCAAAGTGGATCATTCAATAATTCAGACGTAG<br>ACCAAGAGGGGTACTTGAATGTTGCCCAAGTCA<br>ATCAAGACGGTACTAAGTTAAGTTCTGACATCA<br>CGCAAAGTGGTGACTACAATTTCGCCGACGTTG<br>ACCAATCGGGTTCTTTCAATGACTCATCTATCG<br>ACCAAAATGGTACGCAAAATAAGGCAGAGGTA<br>TTGCAATCGGGTCTCAAGAATACGAGTGACATA<br>CACCAAGACGGTAAGCTTAATATAGCCATGGTA<br>GACCAATCTGGGGTTAATAATGACGCCAGTATC<br>GTCCAAGACGGACGTTACCACCAATCAGAGGTC<br>ACTCAATCTGGATACAATGACGTCGCATACTCA<br>TCTCAAAGTGGTTACGAGCACTACTCCTCTATC<br>ACGCAATCAGCATTACCAGGTTTCTTCGGTGGG<br>TTGGCTAGTTTGGCATCCAATAGTGCAACTCAC<br>GTCCAAATGGGAGTTAATAATGCCGCTATCACA<br>ACGCAAGTTGGTGGTGGAAATACAGCTCACGTA<br>TACCAACACTAA |
| **Am18-CsgA** | GVVPQYGGGGNHGGGGNNSGPNMKMKKTKI<br>ASALLLVAATGWASAQEIPASKSVDFNASADF<br>VAAAEGNEITLTQTAPEGNEVGNESVLSQEGD<br>LNAIVVEVTGDANEVLASQMQSGNTFFASVTG<br>NGNFLESEQDSLDSTAEFVVEGDDNLVSLVQL<br>GDGGPFGFIFSRSINSIAGSGNTLSVYQGDGGN<br>WANNAIAGNDNTVFVDQSGDWHESYVRELSG<br>DANLIDVIQDGFYNISDLTVVGSDNEVEVDQD<br>GDENLITWNMVGDGNVLEFEQDGDGNEITTG<br>VFEGSDNEVNIDQIGDVNLATVETIGGGVNTFE<br>INQVGEGNVAYAGVIGLFNEFDLTQVGDANEI<br>STVNFDGWFNTVEIEQDGDANLALAQAGIGD | GGTGTTGTTCCTCAGTACGGCGGCGGCGGTAAC<br>CACGGTGGTGGCGGTAATAATAGCGGCCCAAAT<br>ATGAAGATGAAGAAGACAAAGATCGCAAGTGC<br>ACTCTTGTTAGTCGCTGCTACTGGTTGGGCAAG<br>TGCACAAGAGATACCTGCCTCAAAGTCCGTCGA<br>CTTCAATGCTTCCGCAGACTTCGTCGCCGCTGCC<br>GAGGGTAATGAGATCACATTAACTCAAACTGCT<br>CCCGAGGGAAATGAGGTCGGTAATGAGAGTGT<br>TTTATCCCAAGAGGGAGACTTAAATGCCATCGT<br>TGTCGAGGTCACGGGTGACGCCAATGAGGTATT<br>AGCAAGTCAAATGCAATCTGGTAATACTTTCTT<br>CGCCAGTGTAACTGGGAATGGGAATTTCTTAGA |

| | | |
|---|---|---|
| | AAYVADANLIEMSQEGNENIASVELARDITSS<br>GNEIMVAQTGELNLLDLLVNGNDNVISVMQE<br>GAGNWVTDDMGGQFVISGDMNTFEVTQMGN<br>DNLVTGSITGNGGTVSVTQVGDYNVATVVQM<br>* | GAGTGAGCAAGACTCGTTAGACAGTACGGCAG<br>AGTTCGTTGTAGAGGGAGACGACAATTTGGTTT<br>CCTTAGTTCAATTAGGTGACGGTGGTCCCTTCG<br>GTTTCATCTTCTCCCGTTCGATCAATTCCATCGC<br>TGGTAGTGGGAATACGTTATCAGTATACCAAGG<br>TGACGGTGGTAATTGGGCTAATAATGCTATAGC<br>TGGAAATGACAATACAGTCTTCGTAGACCAAAG<br>TGGAGACTGGCACGAGTCGTACGTTCGTGAGTT<br>GTCCGGTGACGCAAATTTGATCGACGTTATCCA<br>AGACGGGTTCTACAATATCTCAGACCTTACAGT<br>AGTTGGATCTGACAATGAGGTCGAGGTCGACCA<br>AGACGGTGACGAGAATCTCATCACATGGAATAT<br>GGTTGGTGACGGGAATGTATTGGAGTTCGAGCA<br>AGACGGAGACGGTAATGAGATCACGACTGGTG<br>TCTTCGAGGGGTCCGACAATGAGGTTAATATAG<br>ACCAAATAGGGGACGTTAATTTAGCTACTGTTG<br>AGACAATCGGTGGTGGGGTCAATACTTTCGAGA<br>TCAATCAAGTAGGTGAGGGTAATGTAGCATACG<br>CAGGAGTAATAGGGCTCTTCAATGAGTTCGACT<br>TGACGCAAGTTGGAGACGCCAATGAGATCAGT<br>ACAGTTAATTTCGACGGGTGGTTCAATACTGTT<br>GAGATCGAGCAAGACGGTGACGCTAATCTTGCC<br>TTAGCCCAAGCAGGGATCGGTGACGCAGCCTAC<br>GTTGCCGACGCAAATTTAATCGAGATGTCTCAA<br>GAGGGTAATGAGAATATCGCAAGTGTAGAGTT<br>AGCTCGTGACATCACATCCTCAGGTAATGAGAT<br>CATGGTTGCTCAAACAGGTGAGTTAAATTTATT<br>AGACTTGCTTGTTAATGGAAATGACAATGTAAT<br>CAGTGTCATGCAAGAAGGGGCCGGAAATTGGG<br>TTACGGACGACATGGGTGGTCAATTCGTCATAT<br>CCGGAGACATGAATACATTCGAGGTAACTCAAA<br>TGGGGAATGACAATCTTGTTACTGGTAGTATAA<br>CGGGTAATGGTGGTACAGTAAGTGTCACACAAG<br>TCGGTGACTACAATGTCGCCACAGTAGTTCAAA<br>TGTAA |
| **Bc36-CsgA** | GVVPQYGGGGNHGGGGNNSGPNMKTSIYTSV<br>SALALMIGMPAVAQTNTSTVDQTGAAAAATV<br>NQTGSNNTSDIDQNGNGLAYTTGPNTGRTVV<br>ADVEQTGSNGFSQVTQQGGRSSATVDQGGTG<br>MRSTITQGNASTDLGNTATVIQNGTGGGVGTG<br>STITQTGGTGQAYVNQGAATNGAISTITQTGN<br>NQRASVFQTSGAADSSVSQSGGAANVFVSQD<br>GSSESDITQTGGNSEASVRQIGDGNTSLIEQTG<br>VNGDVGDPDNNLANQDTNIGVSQTGDNNSST<br>VRQTGNDQVADVIQTGDFNTASITQDGNFAD<br>ASVTQTGNNNTGTVIRQDGDGSGDNDPASLPV<br>IPTTNATADISQTGDFNEASISQAAVPVEATITQ<br>LGDSNDSSIVQSSTASGAKATNLQTSDNNLSTI<br>TQDSDAEASVTQGGSQFNPGPFAGRDNNVSTV<br>AQGTGAAGSLATVNQDGVLNLSDINQNAANS<br>TASVTQTNIFNTSSVIQSGSGGATADVTQGGG<br>YGDNVSNITQTTNASATVVQTGQPSVGPDFTN<br>ISDIVQNGTGGIASVTQNGSNNESDVAQGGTD<br>GDATVDQDGTFLRSTITQTGGEATALVTQTGN<br>ANVSTISQSAASAGADATVTQSGGENRSDVNQ<br>TGAAKAVVTQAGTEVGYNLAPPNNDSFVVQS<br>ADGADANVSQTGELNLSNVNQSGGASEADVT<br>QTGVYNRSTVTQTVGGAKATVTQSGDPANGG<br>PGVGQGDNLSIVSQSGASTATVTQTANVPTNG<br>FPSNDSFVSQTGDDSDATVDQTGNDNASDVFQ<br>GSDAIADVTQDGTNNTSTVTQSAATSAFVEQI<br>GARNTSTVTQSGGSTVAPYAAYVQQLGNDGL<br>SVVEQDGANNEATLTQSAGSSFADSSIDQSGN | GGTGTTGTTCCTCAGTACGGCGGCGGCGGTAAC<br>CACGGTGGTGGCGGTAATAATAGCGGCCCAAAT<br>ATGAAGACTTCAATCTACACATCTGTCTCCGCC<br>TTGGCCTTAATGATAGGTATGCCCGCAGTCGCT<br>CAAACGAATACATCCACTGTTGACCAAACAGGT<br>GCTGCCGCAGCAGCCACAGTTAATCAAACAGGT<br>TCAAATAATACATCTGACATCGACCAAAATGGA<br>AATGGTCTTGCATACACTACAGGTCCCAATACA<br>GGTCGTACGGTAGTAGCCGACGTCGAGCAAAC<br>AGGGTCGAATGGATTCTCCCAAGTAACGCAACA<br>AGGCGGAAGAAGTTCTGCAACTGTAGACCAAG<br>GTGGTACTGGTATGCGGTCTACAATCACACAAG<br>GAAATGCATCTACTGACCTTGGGAATACGGCTA<br>CGGTCATCCAAAATGGGACGGGTGGTGGTGTCG<br>GTACAGGATCTACGATCACACAAACTGGTGGAA<br>CTGGTCAAGCTTACGTAAATCAAGGGGCAGCTA<br>CGAATGGAGCAATCAGTACGATCACACAAACT<br>GGTAATAATCAACGGGCTTCGGTCTTCCAAACG<br>TCAGGGGCAGCTGACTCCTCTGTTTCCCAATCA<br>GGTGGGGCAGCAAATGTTTTCGTATCCCAAGAC<br>GGTAGTTCGGAGTCAGACATCACACAAACAGGT<br>GGTAATTCAGAGGCTTCAGTCCGTCAAATCGGT<br>GACGGAAATACTTCTCTCATAGAGCAAACGGGA<br>GTCAATGGTGACGTTGGAGACCCAGACAATAAT<br>TTGGCAAATCAAGACACTAATATAGGTGTTTCC<br>CAAACAGGTGACAATAATAGTTCTACGGTTCGT<br>CAAACAGGAAATGACCAAGTCGCTGACGTAAT<br>ACAAACGGGTGACTTCAATACAGCTTCAATAAC<br>ACAAGACGGAAATTTCGCCGACGCAAGTGTCAC |

| | | |
|---|---|---|
| | NNLADVTQGGFDNTSTIVQSGNNNAATVNQS<br>STGNVSDIIQSGNGNSATVTQGGGVL* | TCAAACTGGTAATAATAATACGGGTACAGTTAT<br>CCGGCAAGACGGAGACGGTTCTGGGGACAATG<br>ACCCTGCATCACTTCCTGTTATACCAACAACAA<br>ATGCAACGGCCGACATCAGTCAAACGGGTGACT<br>TCAATGAGGCCTCTATATCCCAAGCAGCCGTAC<br>CCGTTGAGGCAACAATCACACAACTTGGAGACT<br>CCAATGACAGTAGTATCGTCCAATCATCTACTG<br>CCTCAGGTGCTAAGGCAACTAATCTTCAAACGT<br>CTGACAATAATTTATCTACGATAACGCAAGACA<br>GTGACGCAGAGGCATCGGTCACGCAAGGTGGTT<br>CTCAATTCAATCCCGGGCCATTCGCCGGGCGGG<br>ACAATAATGTATCTACGGTAGCCCAAGGAACGG<br>GTGCAGCCGGTTCATTGGCAACAGTAAATCAAG<br>ACGGGGTCTTAAATTTATCAGACATAAATCAGA<br>ATGCTGCAAATTCTACGGCCTCGGTAACTCAAA<br>CAAATATATTCAATACATCGAGTGTAATCCAAT<br>CTGGGTCTGGTGGTGCCACTGCAGACGTTACGC<br>AAGGGGGGGGGTACGGAGACAATGTTTCCAAT<br>ATCACTCAAACAACGAATGCCTCGGCCACTGTC<br>GTCCAAACGGGTCAACCATCTGTCGGACCAGAC<br>TTCACGAATATCTCGGACATCGTACAAAATGGT<br>ACTGGCGGGATAGCATCAGTTACTCAAAATGGA<br>TCTAATAATGAGTCAGACGTTGCCCAAGGTGGT<br>ACAGACGGGGACGCTACGGTAGACCAAGACGG<br>GACATTTCTCCGTTCCACTATCACACAAACTGG<br>CGGAGAGGCTACGGCCTTGGTTACTCAAACGGG<br>TAATGCTAATGTTAGTACGATCAGTCAATCGGC<br>AGCCAGTGCTGGAGCTGACGCCACTGTTACTCA<br>ATCAGGCGGAGAGAATAGATCGGACGTAAATC<br>AAACAGGAGCAGCTAAGGCAGTAGTTACGCAA<br>GCTGGGACTGAGGTCGGTTACAATCTTGCACCC<br>CCGAATAATGACTCCTTCGTCGTTCAATCAGCC<br>GACGGAGCAGACGCCAATGTCTCTCAAACGGGT<br>GAGTTAAATCTTTCGAATGTTAATCAAAGTGGC<br>GGAGCGTCCGAGGCTGACGTAACACAAACAGG<br>TGTATACAATCGTTCGACAGTTACACAAACAGT<br>AGGTGGTGCCAAGGCCACGGTTACACAATCGG<br>GTGACCCTGCTAATGGTGGACCAGGGGTAGGAC<br>AAGGTGACAATTTATCCATCGTCAGTCAATCTG<br>GTGCTTCTACGGCTACAGTTACTCAAACTGCTA<br>ATGTTCCAACGAATGGATTCCCATCCAATGACT<br>CCTTCGTTTCGCAAACGGGAGACGACAGTGACG<br>CTACAGTAGACCAAACGGGAAATGACAATGCA<br>AGTGACGTATTCCAAGGTAGTGACGCCATCGCT<br>GACGTAACGCAAGACGGTACTAATAATACTTCT<br>ACGGTCACGCAATCTGCAGCAACTTCGGCCTTC<br>GTTGAGCAAATAGGGGCCCGTAATACGAGTACT<br>GTCACTCAATCTGGTGGTTCTACAGTTGCGCCTT<br>ACGCAGCTTACGTTCAACAACTCGGTAATGACG<br>GTCTTAGTGTTGTTGAGCAAGACGGGGCCAATA<br>ATGAGGCAACACTTACACAAAGTGCCGGGTCGT<br>CGTTCGCTGACTCGTCTATCGACCAAAGTGGTA<br>ATAATAATCTTGCCGACGTAACTCAAGGTGGAT<br>TCGACAATACAAGTACAATCGTCCAAAGTGGGA<br>ATAATAATGCTGCCACTGTTAATCAAAGTAGTA<br>CAGGGAATGTCTCAGACATAATACAATCTGGGA<br>ATGGAAATTCAGCAACTGTCACACAAGGTGGTG<br>GGGTACTTGAAAACCTGTATTTTCAGGGCGGCT<br>CTGGTGGCTCTGGTGGCTCTGGCGGCAGCGGGC<br>ATCACCACCACCATCATTAA |
| **El43-CsgA** | GVVPQYGGGGNHGGGGNNSGPNMRLWARQT<br>KRSSSAGVLGEILMKKTALLGVSIIALSAAAPA<br>FGQSQSTVTQTGNNSTVDVTQGGPDGGNVST<br>VDQSASDSNVTINQEGFDDNLSGVTNTSTVTQ | GGTGTTGTTCCTCAGTACGGCGGCGGCGGTAAC<br>CACGGTGGTGGCGGTAATAATAGCGGCCCAAAT<br>ATGCGGCTTTGGGCCCGTCAAACTAAGCGTTCT<br>TCTTCAGCAGGTGTCTTAGGTGAGATATTAATG |

| | | |
|---|---|---|
| | AGLNQDVQSTQVGDDQTSSVIQSGVDMEALL<br>VQGGAGNSSTINQSDTDNFADVIQDGDDNISN<br>VSQSDEDGTVDVDQLGERLTSTISQQDDNNQN<br>AVVLQTNADNTSTITQRASNSDVFVTQSGAEN<br>TSTVLQGNISDNQEATILQTGDRNSSDVQQGF<br>DLSGPLDDDNVTFVTQNGDDNTAVVRQINER<br>NLADILQEGDRNEARVVNQGTGVSGGGNEDN<br>IADIDQFGDDNFASINQPGEDGTFTVVQNGFDS<br>SSTGLQSGSNDTASVIQNGEFDSSNVEQSGSGN<br>NATVTQEGFDLDGSLLANSSTVLQSGTGDTAT<br>VSQVGDNQSSFVAQSGSVSTASVVQGGVGNS<br>SGITQSGTDVTATVNQEGDGNSSTVTQANSLS<br>DAVVSQTGDFDTSIVAQSGIDEQANVDQAGET<br>GVSEVTQSGDSNFADVFQRGAENTSFITQSGL<br>DGNVDLDQTGDENFSNAIQAGFEGDIIVNQTG<br>DGNSSIVNQQGDATGFNFATIDQSGNGNTSEAI<br>QFNNSNNANILQSGDDNTSFATQTGTGVIADV<br>DQIGEGATSTVTQDGSGNLAIVSQEGIDFDLSG<br>TTDTSLVAQTGTGQNAEVTQVGDGNSSSVVQ<br>NNADNTASVLTAGIGNSSFVTQNGLTGLATVD<br>QQGDGNSSIVSQGGSNASADVSQQSDTSSSNV<br>NQAGDDVLALVDQAGANHSASIVQLTTGPTG<br>GVDLNNGALIDQQGDGNTANITQSGGIGVFFN<br>VLSGTGLSSGQIAEISQNGSGNTGTVAQSGTTN<br>IGRFFQDGDDNSATISQTGQDSDSSFGQTGNN<br>NTANVTQALIAPPGGIAFADPNQNGDGNSATV<br>VQNGTVTGFFTTRAQSAQIGNGNTLTTTQSGT<br>DDIVFIDQIGDGNMSMVNQLAGGVSNEGDINQ<br>TGDDGISELTQSGTDQFAELFQSGDLNTSLITQ<br>AGVTNTATVTQGSDGNFSSVNQNGTGNSTTV<br>TQ* | AAGAAGACTGCCTTGTTGGGTGTTTCCATAATA<br>GCTCTTTCCGCCGCAGCTCCCGCATTCGGACAA<br>TCACAATCCACTGTCACACAAACAGGAAATAAT<br>TCGACGGTTGACGTAACGCAAGGCGGTCCCGAC<br>GGTGGGAATGTATCCACTGTAGACCAATCAGCT<br>TCCGACAGTAATGTTACTATCAATCAAGAGGGG<br>TTCGACGACAATCTTTCAGGAGTTACTAATACT<br>TCAACTGTCACGCAAGCCGGACTTAATCAAGAC<br>GTTCAATCAACACAAGTTGGGGACGACCAAACT<br>TCGTCGGTAATCCAATCTGGAGTCGACATGGAA<br>GCACTTCTCGTTCAAGGCGGAGCCGGAAATTCT<br>TCCACTATCAATCAATCCGACACAGACAATTTC<br>GCCGACGTCATACAAGACGGTGACGACAATAT<br>ATCTAATGTTTCGCAATCGGACGAGGACGGAAC<br>TGTTGACGTCGACCAATTAGGTGAGCGACTTAC<br>ATCTACTATATCACAACAAGACGACAATAATCA<br>AAATGCCGTTGTACTCCAAACAAATGCAGACAA<br>TACATCCACAATAACACAACGTGCTTCCAATTC<br>AGACGTATTCGTTACACAATCCGGAGCAGAGAA<br>TACTTCCACTGTACTTCAAGGGAATATATCCGA<br>CAATCAAGAGGCAACTATATTACAAACTGGTGA<br>CCGTAATTCTAGTGACGTCCAACAAGGATTCGA<br>CCTTTCGGGACCTTTGGACGACGACAATGTCAC<br>GTTCGTCACGCAAAATGGTGACGACAATACTGC<br>CGTTGTTCGACAAATAAATGAGCGGAATTTAGC<br>CGACATCTTACAAGAGGGAGACCGTAATGAGG<br>CACGTGTCGTCAATCAAGGAACAGGGGTTTCAG<br>GGGGGGGTAATGAGGACAATATAGCCGACATC<br>GACCAATTCGGGGACGACAATTTCGCAAGTATC<br>AATCAACCTGGTGAGGACGGGACTTTCACGGTA<br>GTTCAAAATGGATTCGACTCCTCATCAACAGGT<br>TTACAATCCGGTTCCAATGACACTGCTTCCGTC<br>ATCCAAAATGGGGAGTTCGACTCTTCAAATGTC<br>GAGCAATCCGGATCTGGGAATAATGCTACTGTC<br>ACACAAGAGGGTTTCGACTTAGACGGTTCACTT<br>CTTGCTAATTCCTCTACTGTCTTACAAAGTGGTA<br>CGGGTGACACGGCCACAGTAAGTCAAGTTGGTG<br>ACAATCAATCGTCCTTCGTAGCACAATCGGGTT<br>CGGTATCTACGGCCTCAGTTGTCCAAGGTGGGG<br>TCGGTAATTCTTCTGGTATCACTCAAAGTGGTA<br>CGGACGTAACAGCCACAGTTAATCAAGAGGGT<br>GACGGGAATTCCTCTACTGTCACACAAGCTAAT<br>TCATTATCGGACGCTGTAGTTTCTCAAACGGGA<br>GACTTCGACACAAGTATCGTTGCACAAAGTGGA<br>ATAGACGAGCAAGCAAATGTTGACCAAGCAGG<br>TGAGACAGGAGTCTCCGAGGTAACGCAAAGTG<br>GTGACTCCAATTTCGCCGACGTCTTCCAACGTG<br>GAGCCGAGAATACGAGTTTCATAACACAATCCG<br>GTTTGGACGGGAATGTCGACTTGGACCAAACAG<br>GAGACGAGAATTTCTCCAATGCAATACAAGCTG<br>GTTTCGAGGGTGACATCATAGTCAATCAAACTG<br>GGGACGGTAATTCTTCAATAGTTAATCAACAAG<br>GGGACGCTACTGGTTTCAATTTCGCTACCATAG<br>ACCAATCAGGGAATGGTAATACCTCTGAGGCAA<br>TCCAATTCAATAATAGTAATAATGCTAATATCT<br>TGCAATCAGGTGACGACAATACGTCATTCGCCA<br>CGCAAACGGGTACTGGGGTTATAGCCGACGTCG<br>ACCAAATCGGAGAAGGGGCCACATCTACGGTC<br>ACACAAGACGGAAGTGGAAATCTTGCAATCGTT<br>TCACAAGAGGGTATCGACTTCGACCTTTCAGGA<br>ACTACGGACACTTCACTTGTTGCCCAAACGGGA<br>ACTGGTCAAAATGCCGAGGTCACACAAGTTGGA<br>GACGGTAATTCCTCATCTGTAGTACAAAATAAT<br>GCTGACAATACGGCTAGTGTTTTAACAGCTGGT |

| | | |
|---|---|---|
| | | ATCGGGAATTCATCCTTCGTTACACAAAATGGG<br>CTTACAGGTCTTGCCACTGTTGACCAACAAGGA<br>GACGGTAATTCATCCATCGTTTCACAAGGTGGT<br>TCTAATGCCAGCGCAGACGTAAGTCAACAATCC<br>GACACTAGTTCCTCAAATGTTAATCAAGCTGGT<br>GACGACGTCTTAGCCCTCGTCGACCAAGCAGGT<br>GCAAATCACTCAGCATCTATCGTCCAATTGACA<br>ACAGGACCTACGGGTGGTGTAGACTTAAATAAT<br>GGGGCATTAATAGACCAACAAGGTGACGGAAA<br>TACTGCTAATATCACACAAAGTGGTGGTATCGG<br>TGTCTTCTTCAATGTATTGAGTGGAACTGGGCTT<br>TCTTCAGGTCAAATCGCCGAGATAAGTCAAAAT<br>GGTTCAGGTAATACAGGTACGGTAGCCCAATCT<br>GGGACGACGAATATAGGACGATTCTTCCAAGAC<br>GGAGACGACAATTCCGCCACTATCTCCCAAACT<br>GGACAAGACTCGGACTCGTCTTTCGGACAAACA<br>GGAAATAATAATACGGCTAATGTCACTCAAGCA<br>CTTATCGCTCCGCCCGGCGGAATCGCATTCGCA<br>GACCCAAATCAAAATGGTGACGGGAATTCAGCT<br>ACTGTAGTTCAAAATGGAACGGTCACAGGATTC<br>TTCACGACACGTGCTCAATCCGCCCAAATAGGT<br>AATGGTAATACTCTCACGACGACACAATCTGGT<br>ACTGACGACATAGTTTTCATCGACCAAATCGGT<br>GACGGAAATATGTCCATGGTAAATCAACTTGCT<br>GGCGGAGTTAGTAATGAAGGGGACATAAATCA<br>AACTGGAGACGACGGGATCTCAGAGCTTACGC<br>AATCCGGTACGGACCAATTCGCCGAGCTCTTCC<br>AAAGTGGTGACCTTAATACATCGTTAATAACGC<br>AAGCTGGGGTCACAAATACAGCCACTGTCACAC<br>AAGGTTCCGACGGTAATTTCTCTTCTGTTAATCA<br>AAATGGAACAGGGAATAGTACGACTGTCACGC<br>AAGAAAACCTGTATTTTCAGGGCGGCTCTGGTG<br>GCTCTGGTGGCTCTGGCGGCAGCGGGCATCACC<br>ACCACCATCATTGA |
| **Er46-CsgA** | GVVPQYGGGGNHGGGGNNSGPNMKFSARKT<br>LLLTGVAVLAMSAAASAQEANTGYIGQRDDS<br>NFAYLSQEGANRGFIYQHGQQNVFLPLAAGG<br>AAGDAQQGKNDLGVMQLGSKNVVGGDFLQK<br>GENVGGIAQFGTNNLVDKFRQDGANRGTIFQG<br>GNANVARNTQEEGDSGNDTGIIQLNDSPNKPG<br>NYAESNQLGANGGNFSGILQNGSGNAAINNQT<br>DETSSNAVTAQSGTGNAAINNQTASIDGNAGI<br>LQVGKGNGAINTQTGSSLDATAIIAQNGKDNS<br>AVNTQTNVDPTSATVIQDGNNNAAANVQTSV<br>NNSNAFILQDGNNNTASNLQSPGTQAGPADSL<br>NASIAQSGNKNNAVNLQAGGSGGGKKNTAQT<br>VQDGNRNTALNSQIGDYHNNGYNNDNKAVIG<br>QVGNDNLAVNTQNDGRNSTATILQGGDRNDA<br>YNLSEGGTNHSSAIAQFGDDNLAVNARNRGG<br>NPLEGNDAAILQFGDDNIAANYQNDAIGSEQF<br>TNNSANIGQFGSGNIATNFQDTGSDNEGNITQF<br>GTNNGASNAQVGGGVENTANILQIGSGNSGA<br>NVQSGGTEATANIVQGGSNNVAANLQTNVDP<br>TNASILQLGSRNTAGNVQADLVNSDALIVQDG<br>ARNEASNLQTDGTNMATVIVQKGDRNNAGTI<br>QAGSTDVDAFTLQNGDRNEASNGQSGVTGGT<br>AGIIQDGNRNVAGSVQLTSSDVDASIIQSGNRN<br>DAANLQSGVTGGSAGILQDGRDNTAANLQLS<br>STGVDATIIQSGRFNEAVNSQTASSLSNAFIIQG<br>AGGNNESANVQDDVSGATALIGQFGWGNTAG<br>NLQQADSSNVIAGILQVGSGNTAYNTQGATQT<br>ATYEVLPAFNLTTTASVPGLSNYLTPNMQGPFI<br>FGIAGGPTTLTGSGSAPAEDSIALVAQFGNDNS<br>AANVQIGQLIAPASGIASITTTTYGINTSVQNGG | GGTGTTGTTCCTCAGTACGGCGGCGGCGGTAAC<br>CACGGTGGTGGCGGTAATAATAGCGGCCCAAAT<br>ATGAAATTTAGTGCGCGCAAAACGCTGCTTCTG<br>ACTGGGGTCGCTGTATTAGCCATGTCTGCCGCT<br>GCGTCAGCTCAGGAGGCAAACACGGGCTATATC<br>GGACAACGTGATGACTCAAACTTTGCGTATCTG<br>AGCCAAGAAGGTGCAAACCGCGGTTTCATTTAT<br>CAACATGGGCAACAGAACGTATTTTTGCCACTG<br>GCTGCCGGAGGCGCAGCAGGCGATGCCCAGCA<br>AGGGAAAAACGACTTAGGCGTAATGCAATTAG<br>GTAGTAAAAATGTTGTAGGCGGGGATTTCTTGC<br>AAAAGGGAGAGAATGTTGGAGGTATTGCTCAA<br>TTCGGGACCAACAACCTGGTGGACAAATTTCGT<br>CAAGACGGAGCGAATCGCGGAACAATTTTCCA<br>AGGCGGCAACGCAAACGTCGCACGCAATACGC<br>AGGAGGAGGGTGATTCCGGTAATGACACTGGC<br>ATTATCCAGCTGAACGACTCTCCAAATAAACCA<br>GGGAATTACGCTGAATCCAATCAGCTGGGCGCA<br>AACGGTGGCAATTTCTCAGGCATTCTTCAAAAC<br>GGTTCCGGTAACGCAGCGATTAATAATCAGACC<br>GACGAGACAAGTTCGAATGCAGTGACCGCCCA<br>GAGTGGAACAGGGAACGCTGCCATTAATAATC<br>AAACTGCCTCCATCGACGGTAATGCCGGTATTT<br>TACAGGTTGGTAAGGGCAACGGCGCAATCAAC<br>ACTCAGACGGGGTCAAGTCTTGATGCAACGGCA<br>ATCATTGCTCAGAATGGAAAAGATAACTCCGCG<br>GTGAACACGCAGACTAACGTAGATCCGACAAG<br>TGCAACAGTGATCCAAGATGGGAATAACAATG<br>CAGCTGCAAACGTTCAGACGTCAGTGAACAATT<br>CAAATGCGTTCATTCTGCAAGATGGTAACAACA |

| | | |
|---|---|---|
| | ASWLLGPKTITHSQAQSTQTTSTPFTNPLGEIN<br>VPTVASAAGIVQVGDDNDAFNFQTVAAGSLA<br>ATVQIGDGNGAANVQTATLGAAAAIVQSGNG<br>NTASNLQNGAILSAAGIIQAGHDNVATTTQIAS<br>LSNAALTIQNGKKNVAGTNQSNTVGNTSLIVQ<br>SSNNSLALVDQTGAGGHSSAVVQAGAYSNAW<br>ISQTGSASNSVVLQYGSGSSDADRNYAIVSQG<br>NSGQNSVVIQAGRANVAFVSQN* | ATACAGCAAGCAACCTTCAATCTCCTGGCACAC<br>AGGCCGGTCCCGCGGATAGCTTAAATGCGAGTA<br>TCGCGCAGAGCGGAAACAAAAATAATGCAGTC<br>AACCTGCAAGCTGGTGGCTCAGGAGGTGGGAA<br>AAAGAACACAGCACAAACAGTACAGGACGGTA<br>ACCGCAATACGGCACTGAACTCACAAATCGGA<br>GATTACCACAACAACGGTTACAATAACGATAAC<br>AAAGCTGTCATTGGGCAGGTTGGAAATGACAAT<br>CTAGCTGTAAATACACAAAACGATGGAAGGAA<br>CAGCACCGCTACGATTTTGCAAGGTGGCGATCG<br>TAACGACGCGTATAACCTGAGCGAGGGTGGCA<br>CGAACCACAGCAGCGCGATAGCGCAGTTTGGTG<br>ACGACAACCTGGCAGTGAACGCGCGTAATCGC<br>GGTGGTAACCCGCTGGAAGGTAACGACGCAGC<br>CATTCTCCAATTCGGCGACGATAACATCGCAGC<br>GAACTACCAGAACGACGCGATTGGTAGCGAGC<br>AGTTCACCAACAACTCTGCCAACATCGGCCAAT<br>TCGGTAGCGGTAACATCGCAACCAACTTCCAGG<br>ACACCGGTAGCGATAACGAGGGCAACATCACC<br>CAGTTCGGTACGAACAATGGCGCGTCCAATGCG<br>CAAGTTGGTGGCGGTGTGGAAAATACCGCTAAT<br>ATCCTGCAGATTGGTTCCGGTAATAGCGGTGCG<br>AATGTGCAATCTGGCGGTACGGAAGCGACCGC<br>GAATATCGTGCAAGGTGGGAGCAATAACGTAG<br>CCGCGAACCTTCAAACAAACGTGGATCCGACCA<br>ATGCGTCGATCTTGCAGCTGGGTTCTCGTAATA<br>CCGCCGGTAACGTCCAGGCGGATCTGGTTAATT<br>CTGACGCACTGATTGTGCAGGACGGTGCACGTA<br>ACGAGGCGAGCAATTTGCAGACCGATGGCACC<br>AACATGGCTACGGTCATCGTGCAAAAAGGTGAT<br>CGTAACAACGCCGGCACCATTCAAGCAGGCAG<br>CACCGACGTGGACGCTTTTACTCTGCAAAACGG<br>TGACAGAAATGAGGCTAGCAACGGTCAAAGCG<br>GCGTTACCGGCGGTACTGCGGGTATCATTCAGG<br>ATGGTAACCGCAACGTCGCTGGTTCAGTTCAGT<br>TGACCAGCAGCGATGTTGATGCGTCCATTATCC<br>AAAGCGGAAATCGTAACGATGCCGCAAACCTG<br>CAGTCCGGCGTTACCGGCGGCTCTGCGGGCATT<br>TTACAAGACGGCCGCGATAATACCGCGGCAAA<br>CCTGCAGCTGAGCTCGACTGGCGTTGACGCGAC<br>CATCATCCAGAGCGGCCGTTTTAATGAAGCCGT<br>TAATTCTCAGACTGCCTCCTCACTGTCCAATGCT<br>TTTATCATTCAGGGCGCTGGCGGGAACAATGAA<br>AGCGCTAACGTGCAAGACGACGTGTCAGGGGC<br>GACTGCATTAATCGGGCAATTTGGTTGGGGAAA<br>TACCGCCGGCAATTTGCAACAGGCGGATTCCTC<br>TAATGTTATCGCGGGAATCCTGCAGGTGGGTTC<br>GGGCAATACCGCGTACAATACGCAAGGGGCCA<br>CGCAAACTGCAACGTATGAGGTCCTTCCTGCCT<br>TTAATTTAACGACGACCGCCTCCGTGCCCGGTT<br>TATCCAACTATTTGACGCCGAACATGCAAGGCC<br>CTTTCATCTTTGGAATTGCGGGTGGACCGACAA<br>CCTTGACCGGGTCCGGCTCTGCTCCCGCAGAGG<br>ACAGTATCGCCCTGGTAGCACAATTCGGGAATG<br>ATAACTCTGCTGCCAATGTGCAAATCGGCCAGT<br>TAATCGCACCGGCTAGTGGGATCGCATCCATCA<br>CAACTACAACTTATGGTATTAATACCAGTGTGC<br>AGAACGGAGGGGCGTCTTGGCTTCTTGGACCTA<br>AGACCATCACTCACTCCCAAGCACAAAGCACAC<br>AAACTACCTCCACTCCGTTCACTAACCCCTTGG<br>GCGAAATTAATGTACCTACCGTTGCTTCCGCGG<br>CCGGGATCGTACAAGTTGGCGACGACAATGATG<br>CATTCAATTTCCAGACCGTGGCGGCGGGTAGTC<br>TGGCGGCCACTGTTCAGATTGGTGATGGCAATG |

| | | |
|---|---|---|
| | | GTGCCGCGAACGTACAAACGGCAACGTTAGGG<br>GCTGCAGCCGCAATTGTGCAGTCAGGGAATGGA<br>AATACAGCATCAAACCTGCAGAACGGCGCCATT<br>CTTTCAGCAGCAGGCATTATTCAAGCAGGGCAT<br>GATAACGTCGCGACTACAACCCAGATTGCCAGC<br>TTGAGTAACGCCGCCTTGACCATTCAGAACGGT<br>AAGAAGAATGTGGCAGGCACCAATCAGAGCAA<br>TACGGTCGGAAATACAAGTCTGATTGTCCAGTC<br>TTCCAATAATTCGCTTGCACTTGTCGATCAAAC<br>AGGAGCGGGGGGTCACTCAAGCGCCGTGGTTC<br>AAGCTGGCGCATACAGTAATGCATGGATCAGTC<br>AGACCGGGTCGGCGTCGAACTCAGTGGTTCTTC<br>AATACGGCAGTGGCTCAAGCGACGCGGATCGC<br>AACTATGCTATCGTCTCGCAAGGGAACTCGGGC<br>CAGAACTCCGTAGTGATTCAGGCTGGGCGTGCT<br>AATGTGGCATTCGTGAGTCAGAACTAA |
| **Hs13-CsgA-IronBP** | GVVPQYGGGGNHGGGGNNSGPNMKLKMTV<br>MAAAIAVSSTALANENESNIAQTGFGNDVSVT<br>QLTQGWGNNNSDVTQVGVNNGATVLQQGW<br>RGRNVSDIDQEGMGNNADALQHGTKNSFDLT<br>QSGMFNNSDSQQLGTGNALRVEQEGVKNNAT<br>TYQSGWRNYADISQSGSFNNSDVDQEGYLNV<br>AQVNQDGTKLSSDITQSGDYNFADVDQSGSFN<br>DSSIDQNGTQNKAEVLQSGLKNTSDIHQDGKL<br>NIAMVDQSGVNNDASIVQDGRYHQSEVTQSG<br>YNDVAYSSQSGYEHYSSITQSALPGFFGGLASL<br>ASNSATHVQMGVNNAAITTQVGGGNTAHVY<br>QHGGSGSSGSGGSGGGSGSSGSGGSGGGSGSS<br>GSGGSGRHRHRHRH* | GGTGTTGTTCCTCAGTACGGCGGCGGCGGTAAC<br>CACGGTGGTGGCGGTAATAATAGCGGCCCAAAT<br>ATGAAGTTGAAGATGACGGTTATGGCCGCAGCT<br>ATCGCAGTTTCATCTACTGCCCTTGCAAATGAG<br>AATGAGTCAAATATCGCCCAAACTGGATTCGGT<br>AATGACGTCTCAGTTACACAATTAACGCAAGGT<br>TGGGGCAATAATAATTCAGACGTTACGCAAGTT<br>GGTGTCAATAATGGTGCAACAGTACTCCAACAA<br>GGTTGGAGAGGTCGGAATGTTAGTGACATAGAC<br>CAAGAGGGTATGGGAAATAATGCCGACGCTTTA<br>CAACACGGAACTAAGAATTCATTCGACCTCACT<br>CAAAGTGGAATGTTCAATAATTCTGACAGTCAA<br>CAACTTGGTACAGGTAATGCTTTACGTGTTGAG<br>CAAGAGGGAGTCAAGAATAATGCAACGACATA<br>CCAAAGTGGATGGCGTAATTACGCAGACATAA<br>GTCAAAGTGGATCATTCAATAATTCAGACGTAG<br>ACCAAGAGGGGTACTTGAATGTTGCCCAAGTCA<br>ATCAAGACGGTACTAAGTTAAGTTCTGACATCA<br>CGCAAAGTGGTGACTACAATTTCGCCGACGTTG<br>ACCAATCGGGTTCTTTCAATGACTCATCTATCG<br>ACCAAAATGGTACGCAAAATAAGGCAGAGGTA<br>TTGCAATCGGGTCTCAAGAATACGAGTGACATA<br>CACCAAGACGGTAAGCTTAATATAGCCATGGTA<br>GACCAATCTGGGGTTAATAATGACGCCAGTATC<br>GTCCAAGACGGACGTTACCACCAATCAGAGGTC<br>ACTCAATCTGGATACAATGACGTCGCATACTCA<br>TCTCAAAGTGGTTACGAGCACTACTCCTCTATC<br>ACGCAATCAGCATTACCAGGTTTCTTCGGTGGG<br>TTGGCTAGTTTGGCATCCAATAGTGCAACTCAC<br>GTCCAAATGGGAGTTAATAATGCCGCTATCACA<br>ACGCAAGTTGGTGGTGGAAATACAGCTCACGTA<br>TACCAACACGGTGGATCTGGTAGCAGCGGCTCT<br>GGTGGTTCTGGGGGCGGAAGTGGCTCCTCTGGG<br>AGCGGGGGGTCGGGTGGTGGCTCGGGTTCATCT<br>GGTAGTGGCGGTTCGGGTCGCCATCGCCATCGC<br>CATCGCCATTAA |
| **Hs13-CsgA-IgG-BD** | GVVPQYGGGGNHGGGGNNSGPNMKLKMTV<br>MAAAIAVSSTALANENESNIAQTGFGNDVSVT<br>QLTQGWGNNNSDVTQVGVNNGATVLQQGW<br>RGRNVSDIDQEGMGNNADALQHGTKNSFDLT<br>QSGMFNNSDSQQLGTGNALRVEQEGVKNNAT<br>TYQSGWRNYADISQSGSFNNSDVDQEGYLNV<br>AQVNQDGTKLSSDITQSGDYNFADVDQSGSFN<br>DSSIDQNGTQNKAEVLQSGLKNTSDIHQDGKL<br>NIAMVDQSGVNNDASIVQDGRYHQSEVTQSG<br>YNDVAYSSQSGYEHYSSITQSALPGFFGGLASL<br>ASNSATHVQMGVNNAAITTQVGGGNTAHVY<br>QHGGSGSSGSGGSGGGSGSSGSGGSGGGSGSS | GGTGTTGTTCCTCAGTACGGCGGCGGCGGTAAC<br>CACGGTGGTGGCGGTAATAATAGCGGCCCAAAT<br>ATGAAGTTGAAGATGACGGTTATGGCCGCAGCT<br>ATCGCAGTTTCATCTACTGCCCTTGCAAATGAG<br>AATGAGTCAAATATCGCCCAAACTGGATTCGGT<br>AATGACGTCTCAGTTACACAATTAACGCAAGGT<br>TGGGGCAATAATAATTCAGACGTTACGCAAGTT<br>GGTGTCAATAATGGTGCAACAGTACTCCAACAA<br>GGTTGGAGAGGTCGGAATGTTAGTGACATAGAC<br>CAAGAGGGTATGGGAAATAATGCCGACGCTTTA<br>CAACACGGAACTAAGAATTCATTCGACCTCACT<br>CAAAGTGGAATGTTCAATAATTCTGACAGTCAA |

| | | |
|---|---|---|
| | GSGGSGADNKFNKEQQNAFYEILHLPNLNEEQ<br>RNGFIQSLKDDPSQSANLLAEAKKLNDAQAPK<br>AGGGGADNKFNKEQQNAFYEILHLPNLNEEQ<br>RNGFIQSLKDDPSQSANLLAEAKKLNDAQAPK<br>AGGGGADNKFNKEQQNAFYEILHLPNLNEEQ<br>RNGFIQSLKDDPSQSANLLAEAKKLNDAQAPK<br>A* | CAACTTGGTACAGGTAATGCTTTACGTGTTGAG<br>CAAGAGGGAGTCAAGAATAATGCAACGACATA<br>CCAAAGTGGATGGCGTAATTACGCAGACATAA<br>GTCAAAGTGGATCATTCAATAATTCAGACGTAG<br>ACCAAGAGGGGTACTTGAATGTTGCCCAAGTCA<br>ATCAAGACGGTACTAAGTTAAGTTCTGACATCA<br>CGCAAAGTGGTGACTACAATTTCGCCGACGTTG<br>ACCAATCGGGTTCTTTCAATGACTCATCTATCG<br>ACCAAAATGGTACGCAAAATAAGGCAGAGGTA<br>TTGCAATCGGGTCTCAAGAATACGAGTGACATA<br>CACCAAGACGGTAAGCTTAATATAGCCATGGTA<br>GACCAATCTGGGGTTAATAATGACGCCAGTATC<br>GTCCAAGACGGACGTTACCACCAATCAGAGGTC<br>ACTCAATCTGGATACAATGACGTCGCATACTCA<br>TCTCAAAGTGGTTACGAGCACTACTCCTCTATC<br>ACGCAATCAGCATTACCAGGTTTCTTCGGTGGG<br>TTGGCTAGTTTGGCATCCAATAGTGCAACTCAC<br>GTCCAAATGGGAGTTAATAATGCCGCTATCACA<br>ACGCAAGTTGGTGGTGGAAATACAGCTCACGTA<br>TACCAACACGGTGGTTCTGGCTCGTCTGGATCG<br>GGGGGAAGCGGTGGTGGTTCGGGCTCATCTGGC<br>TCTGGCGGATCAGGGGGTGGTAGTGGATCTTCA<br>GGATCTGGTGGATCCGGCGCGGATAATAAATTC<br>AATAAAGAGCAGCAGAACGCCTTTTACGAGATT<br>CTTCACTTGCCGAATCTTAATGAGGAACAACGT<br>AACGGGTTTATTCAATCACTGAAAGATGACCCC<br>AGCCAGTCGGCAAATCTTCTTGCCGAAGCGAAG<br>AAACTTAACGACGCACAGGCCCCTAAGGCGGG<br>GGGTGGCGGGGCTGATAATAAGTTTAACAAGG<br>AACAACAAAACGCTTTTTATGAGATCTTGCACT<br>TACCAAACTTGAATGAGGAGCAGCGTAATGGGT<br>TCATTCAGAGCTTGAAGGATGATCCTAGCCAAA<br>GCGCTAACTTGTTGGCAGAGGCGAAAAAGTTAA<br>ACGACGCACAAGCCCCAAAGGCGGGCGGAGGG<br>GGAGCTGACAACAAGTTCAACAAAGAGCAGCA<br>AAATGCCTTCTATGAAATTCTGCATTTGCCGAA<br>TTTAAATGAAGAACAGCGCAATGGTTTTATCCA<br>ATCGTTAAAAGACGATCCATCGCAATCCGCCAA<br>CCTTCTGGCGGAAGCGAAGAAGCTGAATGATGC<br>ACAAGCACCGAAAGCGTAA |
